# Conservation genomic analysis reveals ancient introgression and declining levels of genetic diversity in Madagascar’s hibernating dwarf lemurs

**DOI:** 10.1101/620724

**Authors:** Rachel C. Williams, Marina B. Blanco, Jelmer W. Poelstra, Kelsie E. Hunnicutt, Aaron A. Comeault, Anne D. Yoder

## Abstract

Madagascar’s biodiversity is notoriously threatened by deforestation and climate change. Many of these organisms are rare, cryptic, and severely threatened, making population-level sampling unrealistic. Such is the case with Madagascar’s dwarf lemurs (genus *Cheirogaleus*), the only obligate hibernating primate. We here apply comparative genomic approaches to generate the first genome-wide estimates of genetic diversity within dwarf lemurs. We generate a reference genome for the fat-tailed dwarf lemur, *Cheirogaleus medius*, and use this resource to facilitate analyses of high-coverage (~30x) genome sequences for wild-caught individuals representing species: *C. sp. cf. medius, C. major, C. crossleyi* and *C. sibreei*. This study represents the largest contribution to date of novel genomic resources for Madagascar’s lemurs. We find concordant phylogenetic relationships among the four lineages of *Cheirogaleus* across most of the genome, and yet detect a number of discordant genomic regions consistent with ancient admixture. We hypothesized that these regions could have resulted from adaptive introgression related to hibernation, indeed finding that genes associated with hibernation are present, though most significantly, that gene ontology categories relating to transcription are over-represented. We estimate levels of heterozygosity and find particularly low levels in an individual sampled from an isolated population of *C. medius* that we refer to as *C. sp. cf. medius*. Results are consistent with a recent decline in effective population size, which is evident across species. Our study highlights the power of comparative genomic analysis for identifying species and populations of conservation concern, as well as for illuminating possible mechanisms of adaptive phenotypic evolution.

## Introduction

We are in a race against time to preserve species and habitats, and an urgent goal is to identify those populations that are most immediately threatened by shrinking geographic distributions and long-term decreases in population size. Extinction rate estimates reveal recent and rapid loss of biodiversity, with mammals showing the highest extinction rates (Ceballos et al. 2015; Ceballos and Ehrlich 2018). Anthropogenic activity has been identified as a primary driver of this global environmental change (Dirzo et al. 2014), and leads to habitat loss, overexploitation, and accelerated climate change (Haddad et al. 2015). The distributions of non-human primates overlap extensively with a large and rapidly growing human population, and the unintentional battle for resources has resulted in an estimated ~60% of species being threatened with extinction and ~75% of populations in decline (Estrada et al. 2017). The lemurs of Madagascar comprise a clade that contains roughly 20% of all primate species and starkly illustrates the negative impacts of anthropogenic activity on non-human primates (Burns et al. 2016; Salmona et al. 2017): >90% of lemur species are currently threatened with extinction based on International Union for Conservation of Nature (IUCN) Red List assessments (IUCN 2019).

Madagascar’s biota is not only largely endemic but also highly diverse (Goodman and Bernstead 2007). The unique geological and evolutionary history of the island, combined with the contemporary degradation of habitats, has led to an urgent need to understand both their evolutionary history and the ecological interactions between lemurs and the ecosystems in which they occur (Albert-Daviaud et al. 2018). For example, Quaternary climatic variation created periods of glaciation worldwide that resulted in cooler and more arid climates at lower elevations in Madagascar, whereas periods of glacial minima resulted in warmer and more humid climates (Rakotoarinivo et al. 2013). These climatic shifts have been proposed as a driver of speciation in Madagascar (Wilmé et al. 2006), and may have left signatures of population size changes, divergence and introgression in the genomes of the endemic biota. Based on broad climatic trends, one prediction is that low altitude species would have experienced changes in the location and size of suitable habitat, this could have led to changes in N_e_ that may have left a detectable signature in the genome. Similarly, we expect that high-altitude species will have had more stable N_e_ through time. Our study aims to investigate these possibilities.

Here, we focus on Madagascar’s dwarf lemurs (genus *Cheirogaleus*). The genus has undergone substantial taxonomic revisions in recent years (Rumpler et al. 1994; Pastorini et al. 2001; Groeneveld et al. 2009, 2010, 2011, Lei et al. 2014, 2015; Frasier et al. 2016; Mclain et al. 2017), though four species complexes have remained generally well resolved: *Cheirogaleus sibreei*, *C. medius*, *C. crossleyi* and *C. major*. All dwarf lemurs are small (between 150-450 g), nocturnal, obligate hibernating primates. They are distributed across Madagascar and its heterogeneous habitats, from high plateau to low elevation, in the dry west of the island, the tropical east (Goodman and Bernstead 2007), and occur sympatrically in many of these habitats (Blanco et al. 2009). All populations undergo seasonal hibernation despite living in drastically different climatic environments, wet to dry, warm to cold, with some species hibernating for up to seven months of the year. The hibernation phenotype is suspected to be ancestral for the clade, although with subsequent variation in its manifestation (Faherty et al. 2018). For example, *C. sibreei* occupies cold environments at high elevation and hibernates for up to seven months per year (Blanco et al. 2018), while the sister species *C. major* and *C. crossleyi* typically occur at lower altitudes and hibernate for shorter periods of time (approximately four months; Blanco et al. 2018). *C. medius* - displays a similar hibernation phenotype to that of *C. sibreei*, hibernating for up to seven months of the year, but in the highly seasonal, dry deciduous forests of the west.

Conservation efforts must consider current demographics when designing management strategies for populations in decline (Harrison and Larson 2014). Effective population size (N_e_) is a well-established parameter for identifying populations at risk (Nunziata and Weisrock 2018; Yoder et al. 2018), giving insight into the strength of drift and the degree of inbreeding (Chen et al. 2016). Genetic diversity is proportional to N_e_ in a population of constant size (Kimura 1968), and estimates of changes in N_e_ though time can help us better understand potential selective and demographic forces that have affected present day populations (Prado-martinez et al. 2013; Figueiró et al. 2017; Árnason et al. 2018; Vijay et al. 2018). Failure to accommodate for changes in genetic diversity over time in populations of conservation concern can lead to elevated extinction risks (Pauls et al. 2013) given that levels of quantitative genetic variation necessary for adaptive evolution become reduced, and deleterious mutations can accumulate (Rogers and Slatkin 2017). Though N_e_ is typically considered at the population level, it is also important to assess how diversity varies among species. Characterizing genetic diversity in a phylogenetic context requires an informed understanding of species boundaries, and the very nature of species boundaries carries the expectation that phenotypes and genomic regions remain differentiated in the face of potential hybridization and introgression (Harrison and Larson 2014).

By investigating the ancestral demography of living species we can make inferences about their current genetic health and prospects for survival (Prado-martinez et al. 2013). The development of methods such as pairwise and multiple sequentially Markovian coalescent (PSMC and MSMC) allows for the use of a single biological sample to give insight into an individual’s ancestry, and theoretically, to the changes in conspecific N_e_ over considerable time periods (Li and Durbin 2011; Schiffels and Durbin 2014). These techniques have been used to identify specific timescales of species declines, such as in the case of the woolly mammoth (*Mammuthus primigenius*) wherein genomic meltdown in response to low N_e_ is thought to have contributed to its extinction (Palkopoulou et al. 2015; Rogers and Slatkin 2017). Understanding the speed of ancestral population declines, from the reduction in N_e_ and the accumulation of detrimental mutations, to genomic meltdown, is critical for the conservation of extant species. Importantly, effective population size, genetic diversity, and species boundaries can all be used in conservation planning and, directly used to calculate IUCN “Red List” qualification.

A genomic approach offers increased power and precision to addressing the questions outlined above. For example, by looking at patterns of genetic variation found within hominid genomes, numerous studies have revealed instances of interbreeding and adaptive introgression that occurred between early modern humans and archaic hominids (Hajdinjak et al. 2018; Slon et al. 2018). Admixture between Neanderthals and modern humans has, for example, left detectable signatures in the genome sequences of modern humans (Harris and Nielsen 2016), which can be identified in the genome sequences of single individuals (Green et al. 2010a; Hajdinjak et al. 2018; Slon et al. 2018). A focus on making use of the entirety of information contained within a single genome has also been beneficial for a number of non-human study systems where the ability to collect and sequence many samples is prohibitively difficult (Prado-martinez et al. 2013; Meyer et al. 2015; Palkopoulou et al. 2015; Abascal et al. 2016; Figueiró et al. 2017; Rogers and Slatkin 2017; Árnason et al. 2018).

Using whole genome sequences for five individuals across four species of dwarf lemur, we ask if previously published phylogenetic relationships, based on mitochondrial and morphological comparisons (Groeneveld et al. 2009, 2010; Thiele et al. 2013; Lei et al. 2014), are consistent across the genome. We show that despite an overall congruence, there are small regions of introgression, a proportion of which is between *C. sibreei* and *C. sp. cf. medius*, two species with the most similar hibernation profiles. Since dwarf lemurs are the only obligate hibernating primates, and hibernation is thought to be ancestral (Blanco et al. 2018), we tested if genes associated with the hibernation phenotype are present in introgressed regions of the genome. We find evidence that genes implicated in the control of circadian rhythm and feeding regulation (such as NPY2R and HCRTR2) have introgressed between species of dwarf lemur.

Together, this work offers the largest contribution of novel genomic resources for any study to date of Madagascar’s lemurs. These data allow us to better understand contemporary levels of diversity in isolated populations and the demographic history of *Cheirogaleus*. Our approach represents a concerted effort to utilize genomic data for the purposes of formulating hypotheses of conservation threat and to better understand the evolutionary history of this unique clade of primates, as well as future prospects for its survival.

## Methods

### Genome sequencing and assembly

To allow us to compare inter-clade diversity on a genomic scale, we first generated a reference genome from a Duke Lemur Center *Cheirogaleus medius* individual, DLC-3619F, which died of natural causes. DNA was extracted from liver tissue, and shotgun and Chicago libraries prepared by Dovetail Genomics. Each library was sequenced on two lanes of Illumina HiSeq 4000 at Duke Sequencing Core (2×150bp reads; Supplemental Table 1). A *de novo* assembly was constructed using a Dovetail’s Meraculous assembler, and the HiRise software pipeline was used to scaffold the assembly (Putnam et al. 2016). Adapters were removed using TRIMMOMATIC (Bolger et al. 2014), which also discarded reads < 23 bp and *q* < 20. We assessed the completeness of the assembly using standard summary statistics (e.g. number of scaffolds and scaffold N50) and the presence of orthologous genes sequences (BUSCO pipeline; v3; Waterhouse et al. 2017).

To facilitate tests of phylogenetic relationships across the genome, estimate demographic history, and measure genetic diversity, we sampled four wild-caught dwarf lemurs, one each from the species *Cheirogaleus sp. cf. medius, C. major, C. crossleyi*, and *C. sibreei*. Our aim was to include one sample from each of the four primary clades found within *Cheirogaleus* (Groeneveld et al. 2010). Individuals were live-trapped (sampling locations shown in Fig. 1A), ear clips taken, and the animals released. *C. sp. cf. medius* was sampled from a population in Tsihomanaomby, a forest in the north-east of Madagascar outside of the known *C. medius* range, and where *C. sp. cf. medius, C. major* and *C. crossleyi* are found in sympatry. Samples from the *C. sp. cf. medius* population in Tsihomanaomby are distinct from *C. medius* in mtDNA (Fig. 1B) and morphology (Groeneveld et al. 2009). This lineage warrants further investigation and is here referred to as *C. sp. cf. medius*, but was not sampled in the taxonomic revision of *Cheirogaleus* by Lei *et al* (2014). The *C. major* individual was sampled from Marojejy National Park, a rainforest ~45 km from the *C. sp. cf. medius* sampling site, which *C. crossleyi* and *C. sibreei* also inhabit, though the latter is only found at elevations above 1500 m. Sampling sites for *C. sp. cf. medius* and *C. major* are separated by ~45 km. We sampled *C. crossleyi* from Andasivodihazo, Tsinjoarivo (where *C. crossleyi* and *C. sibreei* occur sympatrically) and *C. sibreei* from Ankadivory, Tsinjoarivo. Andasivodihazo and Ankadivory sampling sites are separated by ~6 km but are part of the same continuous forest (Fig. 1A).

**Figure 1.**
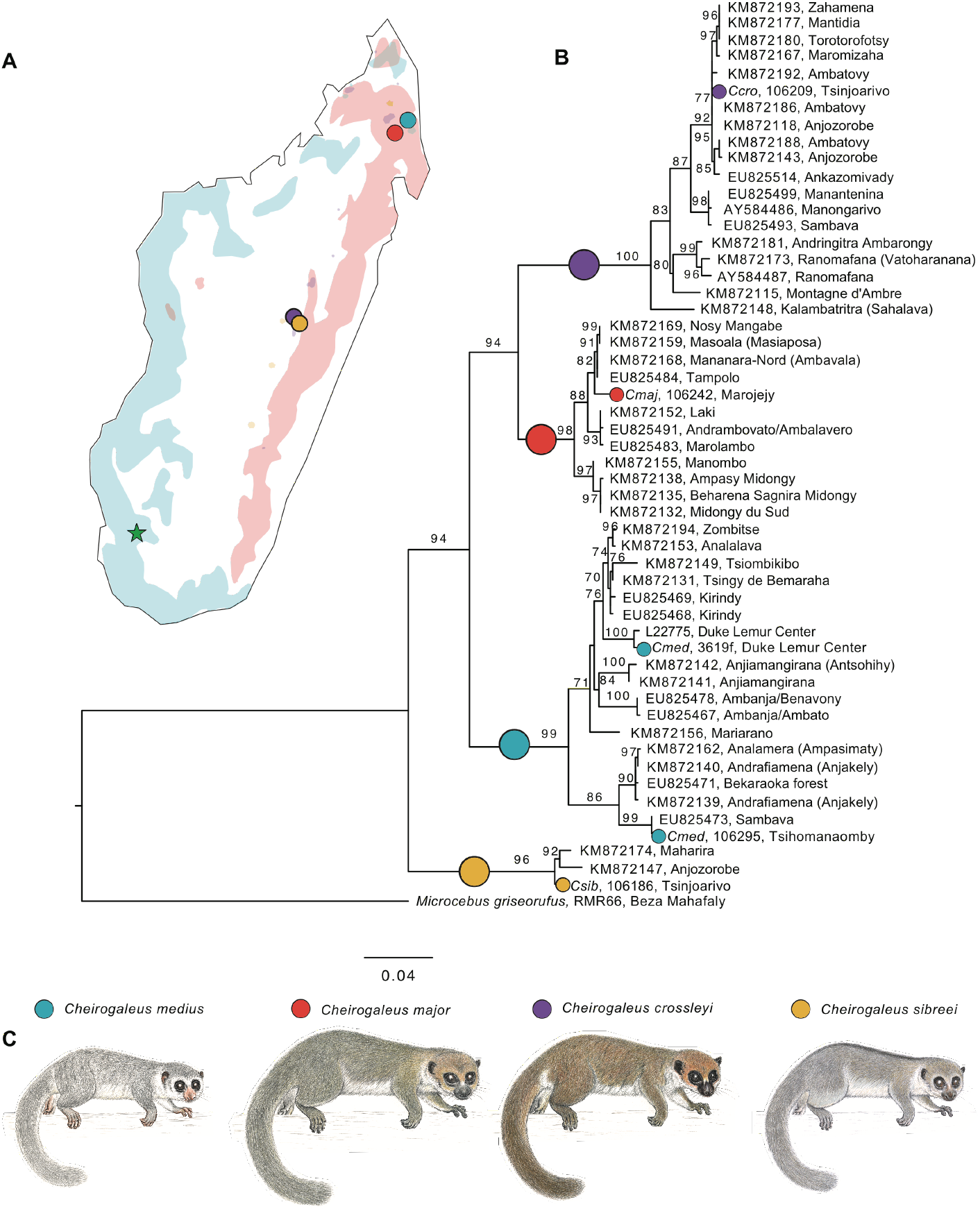
Geographic distributions and phylogenetic relationships of *Cheirogaleus*. **A**. Map of Madagascar showing the sampling locations and IUCN Red List distributions of the four study species. Green star shows sampling location of outgroup *Microcebus griseorufus* **B**. Maximum likelihood estimation showing relationships within *Cheirogaleus* based on 684 bp cytochrome c oxidase subunit II (COII). Individuals used here in genomic analyses are indicated by their species color. Comparative taxa were taken from the National Center for Biotechnology Information (NCBI) online database. Locations for these comparative samples were taken from NCBI or, if unlisted, their publication. Bootstrap values shown. **C.** *Cheirogaleus* illustrations, copyright 2013 Stephen D. Nash / IUCN SSC Primate Specialist Group; used with permission

DNA was extracted using DNeasy (*C. sibreei*) or MagAttract (remaining samples) kits. One library was prepared per individual and barcoded libraries were then pooled prior to sequencing. Sequences were generated using the Illumina HiSeq 4000 machine at Duke Sequencing Core (2 × 150 bp reads), and read quality was assessed using FASTQC and then filtered using TRIMMOMATIC. Reads < 50 bp were discarded after trimming for quality (*q* < 21, leading: 20, trailing: 20). Expected genome size for each of the four species we sampled was estimated using JELLYFISH (Marçais and Kingsford 2011) and FINDGSE (Sun et al. 2018). Reads were aligned to the *C. medius* reference genome using BWA mem (Li and Durbin 2009) with default settings. Bam files were produced using SAMTOOLS (Li et al. 2009) and duplicates were marked using PICARD (Broad Institute, accessed 2018). Reads were then realigned around indels using GATK’s ‘RealignerTargetCreator’ and ‘IndelRealigner’ tools (Mckenna et al. 2010; Depristo et al. 2011). We then genotyped each individual at SNPs mapping to scaffolds >100 kb (99.6% of the assembly). Individual .gvcf files were first generated for each individual using GATK’s ‘HaplotypeCaller’ tool. Joint genotyping was then carried out for all samples using GATK’s ‘GenotypeGVCFs’ tool. The final set of variants were filtered based on GATK Best Practices (Van der Auwera et al. 2013). One *Microcebus griseorufus* individual (RMR66), sampled from Beza Mahafaly Reserve, was sequenced as approximately 400 bp insert libraries on an Illumina HiSeq 3000 (2 × 150 bp reads) to ~40X and included in the pipeline described above so that it could be used as an outgroup.

### Phylogenetic relationships

To investigate species boundaries between our samples, we first confirmed the species assignment of our individuals using mtDNA barcoding. We used cytochrome c oxidase subunit II (*COII*) sequencing and NCBI Genbank to access a subset of *COII* sequences from across the *Cheirogaleus* clade. Additionally, we used BLAST+ 2.7.1 (Altschul et al. 1990) to extract the *COII* sequences from our genome-wide *Cheirogaleus* data. Sequences were aligned using MAFFT v7.017 (Katoh et al. 2002), totaling 54 sequences with 684 nucleotide sites. We estimated a maximum likelihood (ML) phylogenetic tree for the aligned sequences using ModelFinder (Kalyaanamoorthy et al. 2017), as implemented in IQ-TREE v.1.6.6 (Nguyen et al. 2015), to assess relative model fit. The phylogeny was rooted on the outgroup *Microcebus griseorufus* and significance was assessed using 500,000 ultrafast bootstrap replicates (Hoang et al. 2018).

To assess variation in relationships among species across the genome, we used SAGUARO (Zamani et al. 2013). SAGUARO generates similarity matrices for each region that represents a new relationship, therefore creating statistical local phylogenies for each region of the genome. We used 105,461,324 SNPs, averaging ~four SNPs per 100 bp. Our final set of variants for the four ingroup species was run in SAGUARO using default iterations. We visualised the distribution of relationships across the genome in R (R Core Team 2015), plotting phylogenetic relationships as neighbour joining trees. Topologies that do not agree with the previously accepted taxonomic relationships within the genus (e.g. Thiele et al. 2013; Lei et al. 2014) are from here termed ‘discordant’ trees. Since the *C. medius* reference genome is not annotated, we identified genes in discordant regions by using BLAST+ to infer gene function from homologous genes from the grey mouse lemur (*Microcebus murinus)* NCBI protein database (Larsen et al. 2017). *M. murinus* is the highest quality genome assembly available within the sister clade to *Cheirogaleus*, and is approximately 27 My divergent (Dos Reis et al. 2018).

### Test for independence

Because SAGUARO identified a small number of discordant genomic regions, we next carried out formal tests for admixture between taxa using Patterson’s *D*-statistics (Green et al. 2010b; Durand et al. 2011), as implemented in ANGSD (Korneliussen et al. 2014). We ran two tests; the first test was based on discordant relationships identified in the SAGUARO analyses, and looked for admixture between *C. crossleyi, C. major* and *C. sp. cf. medius*, using *C. sibreei* as the outgroup (O), given the relationship (((P1,P2),P3),O). The second test looked for introgression between *C. sp. cf. medius* (P2) and *C. sibreei* (P3) with outgroup *Microcebus griseorufus*, and using either *C. crossleyi* or *C. major* as P1 (allowing us to test for introgression between different combinations of species).

Patterson’s *D* gives a genome wide estimate of admixture and was not designed to quantify introgression at specific loci (Martin et al. 2015). To identify specific regions of admixture, we used *f*_*d*_ (Martin et al. 2015), which provides a point estimate of the admixture proportion at a locus, allowing for bidirectional introgression on a site-by-site basis. We ran *f*_*d*_ on combinations that had returned a significant Patterson’s *D*, first between *C. sp. cf. medius* and *C. major (C. med/C. maj)*, and secondly between *C. sp. cf. medius* and *C. sibreei (C. med/C. sib)*. We used 40 kb windows with a 10 kb step size.

We defined candidate regions of introgression as those that fell into the top 0.05% of the genome wide distribution of *f*_*d*_. These ‘significant’ regions were extracted and BEDTOOLS (Quinlan and Hall 2010) was used to merge overlapping windows. Windows that overran the length of a scaffold were corrected, reducing the regions of interest from 2.25 Mb to 2.24 Mb. Scaffolds containing windows with significant *f*_*d*_ were filtered for coverage, identifying sites higher than twice the mean, and lower than half the mean coverage in order to match MSMC2 analyses. This mean was averaged across samples on a per site basis. Windows were masked if more than 30% of the sites within a window did not meet filtering criteria.

Given that the hibernation phenotype is thought to be ancestral for the dwarf lemur clade (Blanco et al. 2018), we set out to test the hypothesis that genes associated with hibernation were present in introgressed regions. Genes present in introgressed regions were identified from the *M. murinus* protein database using tblastn, and filtered by evalue(0.0001), max_hsps(1) and max_target_seqs(20). Results were parsed by bit score then evalue to retain only unique RefSeqIDs. Genes associated with metabolic switches before and during hibernation were pulled from the hibernation literature (Faherty et al. 2016, 2018; Grabek et al. 2017) that we refer to as hibernation-associated genes. These genes were chosen from the literature on small hibernating mammals and included all gene expression work in *Cheirogaleus*. Ensembl biomaRt (Durinck et al. 2005, 2009) was used to convert between different terms of reference. The hibernation-associated genes were cross-referenced with genes identified in windows that contained significant *f*_*d*_. To identify over represented biological pathways of introgressed genes, we ran Gene Ontology (GO) enrichment analysis using the R/bioconductor package goseq v.1.32.0 (Young et al. 2010).

### Age of introgression

To assess the timing of introgression, we used the software HYBRIDCHECK v1.0 (Ward and Van Oosterhout 2016). HYBRIDCHECK differs from *f*_*d*_ by using spatial patterns in sequence similarity between three sequences, as opposed to taking phylogeny into account by using a fourth sequences as the outgroup. Additionally, the analysis does not use pre-defined windows. As input for HYBRIDCHECK, we created full-sequence fasta files for each individual. To do so, we first ran GATK v3.8 FastaAlternateReferenceMaker for each individual, replacing bases in the *C. medius* reference genome that were called as non-reference alleles for that individual, using the unfiltered VCF file (see genotyping, above). Next, the following classes of bases were masked: (1) Non-reference bases that did not pass filtering (DP<5, DP>2*mean-DP, qual>30, QD<2, FS>60, MQ>40, MQRankSum<−12.5, ReadPosRankSum<−8, ABHet <0.2 or >0.8); (2) sites that were classified as non-callable using GATK v3.8 CallableLoci (using a minimum DP of 3). HYBRIDCHECK was run for all scaffolds larger than 100kb using default settings, testing all possible triplets for the four *Cheirogaleus* species. We only retained blocks that contained at least 10 SNPs and had a p-value <1×10^−6^. Blocks identified between sister species in any given triplet were removed to reduce the chance of identifying blocks due to incomplete lineage sorting. Estimated dates for introgressed blocks were converted using a per-year mutation rate of 0.2×10^−8^, based on a per-generation mutation rate of 0.8×10^−8^ (Yoder et al. 2016). This mutation rate is currently the most accurate estimate available, calculated for mouse lemurs *(Microcebus murinus)* from average estimates between humans and mice. As the phylogenetically closest estimate for dwarf lemurs, this was used throughout the analyses. We checked for overlap between regions identified using *f*_*d*_ and those identified using HYBRIDCHECK.

### Historical and contemporary effective population size

To estimate how N_e_ has varied through time, for each species, we estimated N_e_ using the software MSMC2 (Schiffels and Durbin 2014; Malaspinas et al. 2016). Due to only sequencing one individual per species, we ran each species as a single population of *n=1* (with a single individual per species, we could not run MSMC2 to estimate cross-coalescence rates between populations). For this analysis, we included scaffolds larger than 10 Mb, using a total of 52 scaffolds, comprising ~2 Gb and 89.23% of the genome assembly (total assembly is 2.2 Gb). SAMTOOLS and a custom script from the MSMC tools repository were used to generate input files per scaffold: a VCF and a mask file to indicate regions of sufficient coverage. Additionally, a mappability mask was generated using SNPABLE REGIONS (Li 2009), providing all regions on which short sequencing reads could be uniquely mapped. To run MSMC2, we used “1*2+30*1+1*2+1*3” to define the time segment patterning. Coalescent-scaled units were converted to biological units using a generation time of four years and a per-generation mutation rate of 0.8 × 10^−8^. To estimate uncertainty in estimates of N_e_, we performed 50 bootstrap replicates. We calculated the harmonic mean of the N_e_ for each individual, excluding both the first five and the last five time segments

To study contemporary patterns of genetic diversity, we ran a sliding window analysis in 100 kb windows across the genomes. As a measure of heterozygosity, we identified total heterozygous sites per window, and sites with missing genotype calls in one or more species. We then divided the number of heterozygous sites by the window size, minus the number of missing genotype calls. We note that this measure differs from heterozygosity in the population-genetic sense (H = 2pq), but is still a useful summary of the genetic variation contained within the populations from which our samples were collected.

## Results

### Genome assembly and sequencing statistics

The final assembly of the Dovetail-generated reference genome for *Cheirogaleus medius*, based on ~110 X coverage, comprised 191 scaffolds >100 kb and had a scaffold N50 of 50.63 Mb. The genome contained 92.7% complete single-copy BUSCO orthologs, 3.4% were fragmented, and 3.2% were missing. For the four wild-caught individuals, Illumina sequencing produced 350.4 Gb of raw data, totaling 1,511,287,743 paired-end reads and representing ~30 X coverage per individual. After quality filtering, 1,444,867,212 (97%) high quality reads were retained (82.1% paired, 12.6% forward, 1.7% reverse) and mapped to the *C. medius* reference genome. Genome size was estimated to be 2.4 Gb for *C. sp. cf. medius*, 2.6 Gb for both *C. major* and *C. crossleyi*, and 2.5 Gb for *C. sibreei*.

### Phylogenetic relationships

ModelFinder within IQ-TREE selected model TIM2+F+G4 for the ML analyses based on the Bayesian Information Criterion and corroborated the overall topology as previously published, confirming the clade-level assignment of the samples we use for re-sequencing in this study (Fig. 1B). The *C. sp. cf. medius* individual phylogenetically groups with individuals identified as *C. medius* using mitochondrial DNA (Fig. 1B), and this specific sub-clade of *C. medius* was not sampled by Lei *et al* (2014). Our *C. major* individual phylogenetically groups into the ‘major C clade’ of Lei *et al.* (2014), and *C. crossleyi* phylogenetically groups with the ‘crossleyi B clade’ of Lei *et al.* (2014). *C. sibreei* remains a single species.

The mitochondrial sequences for the four individuals of *C. sp. cf. medius, C. major, C. crossleyi* and *C. sibreei* used in this study are highly divergent, with species separated in well-defined clades. By using a phylogenomic approach, we were able to confirm the relationships identified using mitochondrial data at a genome-wide scale. To do this, we used SAGUARO to identify the most common relationships among *C. sp. cf. medius, major, crossleyi* and *sibreei* (the *C. medius* reference genome was not included in subsequent analyses due to admixed ancestry (Williams *et al.*, unpublished data). SAGUARO identified 16 unique relationships across the genomes (Fig. 2). We concatenated results that showed the same topology, and from these remaining distance matrices, seven (99.4% of the genome) recapitulate phylogenetic relationships identified using mtDNA. Thus, the vast majority of genomic regions support previously published relationships among dwarf lemur species, which were mostly based on mitchondrial data and morphological comparisons (Groeneveld et al. 2009, 2010; Thiele et al. 2013; Lei et al. 2014). Our SAGUARO results indicate that *C. major* and *C. crossleyi* are genomically most similar to each other (sister species cannot be formally inferred with this method, as trees are unrooted), and equally distant from *C. sp. cf. medius*. *C. sibreei* shows the greatest divergence within the clade, which is in agreement with previous phylogenetic studies. These results thus provide the first genome-wide support for four distinct species of *Cheirogaleus*.

**Figure 2.**
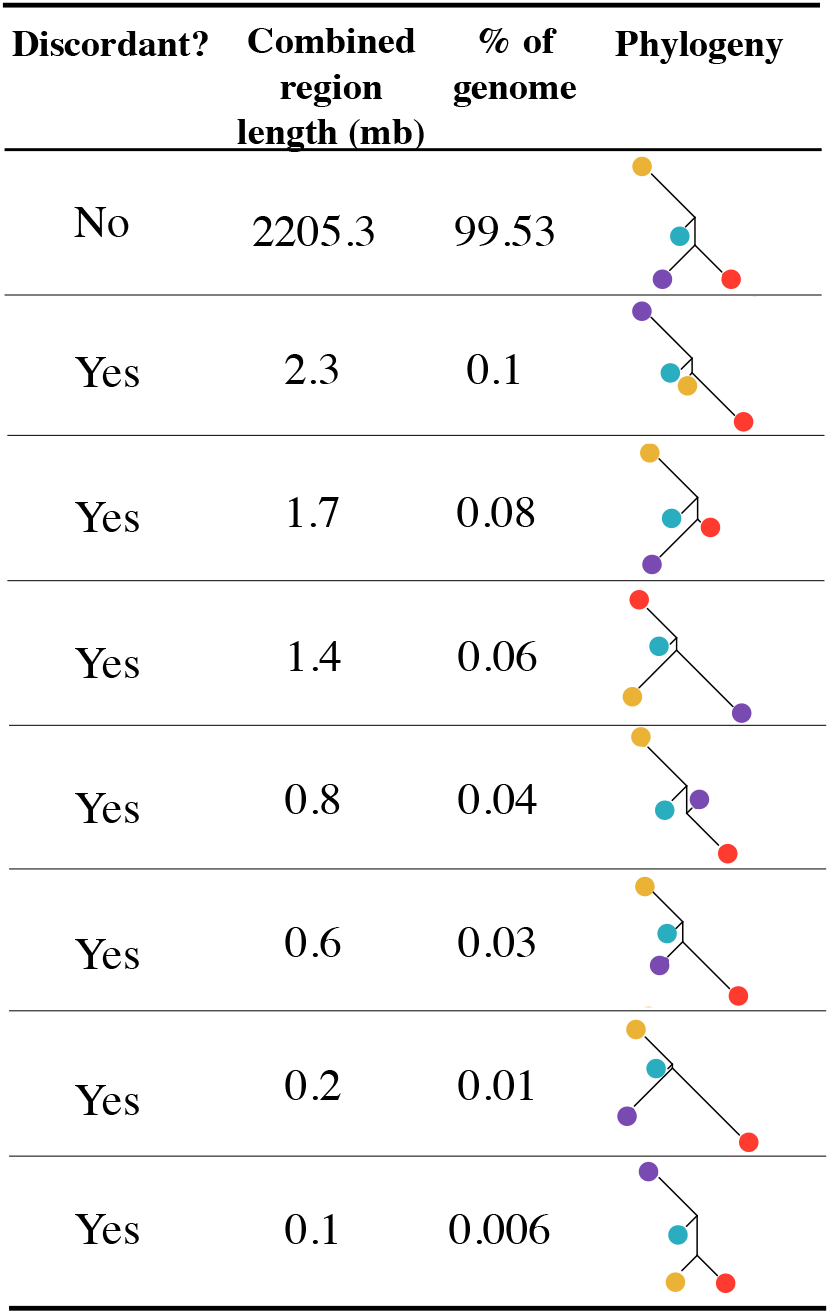
The relationships assigned to regions of the genome that were identified in the SAGUARO analyses. Discordance to previously published phylogenies indicated. Phylogenies are neighbor joining trees generated from the distance matrices. Phylogenies with the same overall relationships have been concatenated. Colors represent species: *Cheirogaleus sp. cf. medius* (blue), *C. major* (red), *C. crossleyi* (purple), *C*.

Additional topologies identified by SAGUARO (Fig. 2) illustrate variation in local phylogenetic relationships across the genome. Therefore, despite the overall support for the species tree, local variation in phylogenetic signal might be driven by evolutionary processes such as variation in mutation rate, incomplete lineage sorting, or introgression. For example, SAGUARO identified seven discordant topologies of which two, spanning 3 Mb of the genome in total, show a close relationship among *C. major* and *C. sp. cf. medius*. Two trees show a topology in which *C. sibreei* and *C. sp. cf. medius* were more closely related (657 kb of the genome), and three trees grouped *C. crossleyi* and *C. sp. cf. medius* as more closely related (2.4 Mb of the genome). SAGUARO is not forced to assign genomic regions to all hypotheses and so two of the 16 relationships were not assigned to regions of the genome. Genes present in discordant regions were identified using tblastn, resulting in a total of 30 unique genes (supp. table 3).

### Signals of ancient introgression

To test if discordant trees were a result of admixture between species, as opposed to incomplete lineage sorting, we used Patterson’s *D*-statistics. When testing for introgression between *C. sp. cf. medius* with either *C. crossleyi* or *C. major*, we found a genome wide excess of ABBA sites, giving a significantly positive Patterson’s *D* of 0.014 ± 0.0008 (*Z* score 16.94) between *C. sp. cf. medius* and *C. major*, thus providing evidence for admixture between *C. major* and *C. sp. cf. medius*. This result is also in agreement with the most common discordant topology identified in the SAGUARO analysis summarized above. When testing for introgression between *C. sp. cf. medius* and *C. sibreei*, we again found a genome wide excess of ABBA sites. With *C. crossleyi* as P1, there was a significantly positive Patterson’s *D* of 0.06 ± 0.001 (Z score 54.37). With *C. major* as P1, there was a significantly positive Patterson’s *D* of 0.06 ± 0.01 (Z score 53.89). Both results are evidence for introgression between *C. sp. cf. medius* and *C. sibreei*.

To identify specific regions of admixture, we used the *f*_*d*_ statistic (Martin et al. 2015; Fig. 3). We calculated *f*_*d*_ for comparisons that had significant genome-wide estimates of Patterson’s *D*: between *C. sp. cf. medius* and *C. major (C. med/C. maj)*, and between *C. sp. cf. medius* and *C. sibreei (C. med/ C. sib)*. We define a region as being introgressed if *f*_*d*_ was in the top 0.05% of the genome wide distribution. The *C. med/C. maj* test (Fig. 3A) resulted in an average of 1,903 biallelic SNPs per window, a mean *f*_*d*_ of 0.0008 and a 0.05% cutoff of 0.137. There were 21 scaffolds containing one or more regions of significant *f*_*d*_. The scaffold with the highest average *f*_*d*_ was ScMzzfj_1392_HRSCAF_67928 (*f*_d_ =0.01). The *C. med/C. sib* test (Fig. 3B) averaged 3,005 biallelic SNPs per window, with an average *f*_*d*_ of 0.01 and a 0.05% cutoff of 0.188. There were 23 scaffolds with regions of one or more significant *f*_*d*_ window, and scaffold ScMzzfj_2194_HRSCAF_93304 had the highest average (*f*_*d*_ =0.07).

**Figure 3.**
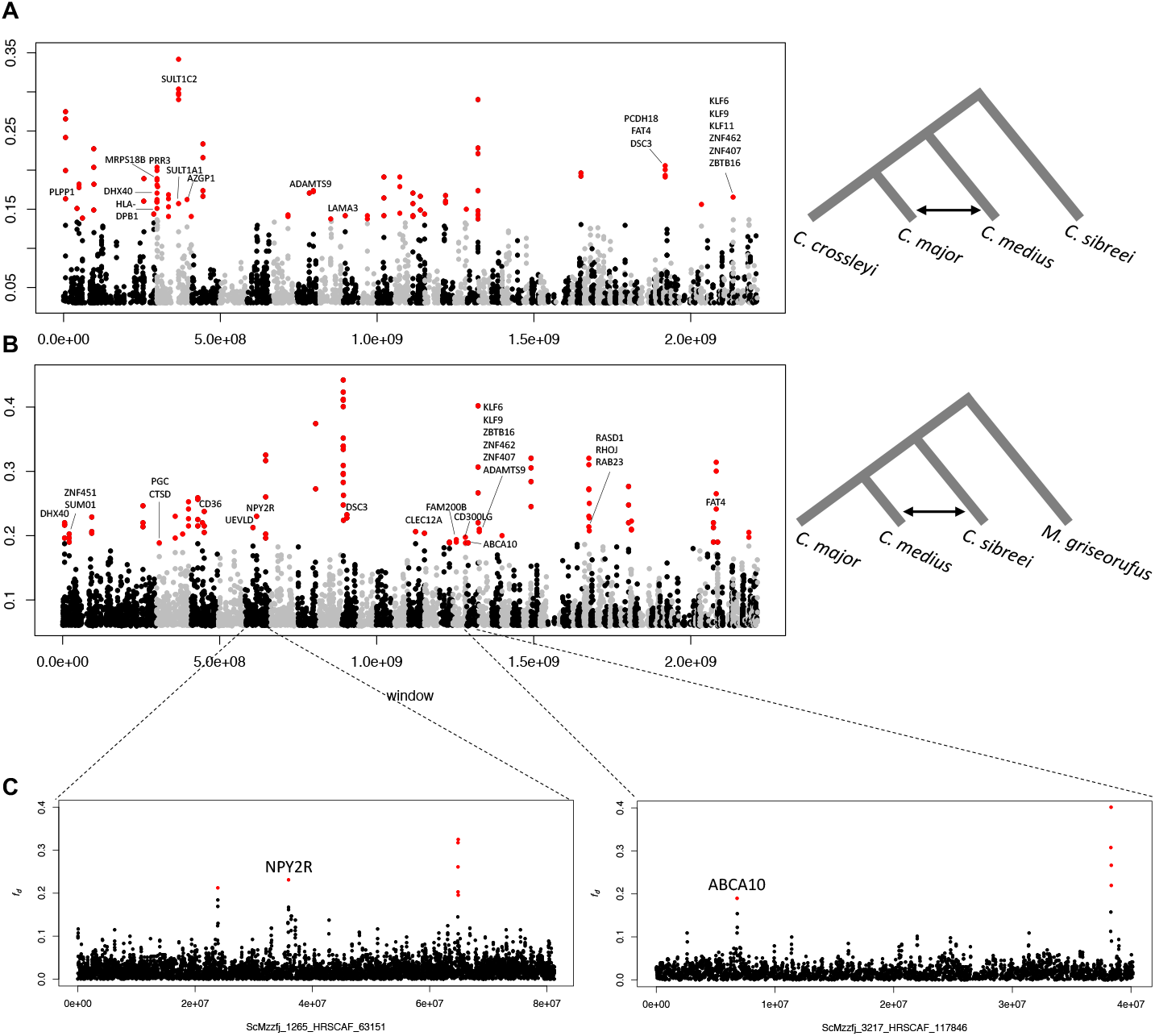
*F*_*d*_ test for introgression plotted across the genome. *F*_*d*_ statistic computed in 40 kb windows with a 10 kb sliding window, with the full phylogeny and associated test indicated on the right. Scaffolds are arranged from largest to smallest, alternating in color. “Significant regions” are shown in red. Hibernation-related genes are labelled. **A.** Results from a test of introgression between *Cheirogaleus sp. cf. medius* and *C. major* (C. med/C. maj). **B.** Results from a test of introgression between *C. sp. cf. medius* and *C. sibreei* (C. med/C. sib) **C.** Isolated scaffolds of interest.

We next used HYBRIDCHECK to estimate the date of the introgression within the clade (Fig. 4). The average age of introgressed regions across all species pairs was 4.12 My, and the average length of introgressed regions was 2.2 kb (Table 1), showing that introgression was predominantly ancient. The lower boundary estimate of the most recent split within the *Cheirogaleus* clade is approximately 6 Mya (identified in Dos Reis et al. (2018) between *C. medius* and *C. major)* and therefore approaches, but does not overlap with, our oldest average estimate of admixture, which is between *C. major* and *C. sibreei* and dated 4.52 Mya. Following the removal of sister species in any given triplet, the number of introgressed regions were calculated between pairs from any given triplet. The largest number of introgressed regions (totaling 9,553) were detected between *C. sp. cf. medius* and *C. sibreei*, generated from testing triplet *‘med/cro/sib’* and *‘med/maj/sib’*. These values were followed by regions detected between *C. sp. cf. medius* and *C. major* (4,959). These results corroborate our *f*_*d*_ findings. Calculating the overlap between *f*_*d*_ and HYBRIDCHECK blocks for *C. med/C. maj* showed that 8 of the 86 *f*_*d*_ blocks overlap with the 4959 HYBRIDCHECK blocks to create a total of 12 discrete overlapping sections. For *C. med/C. sib*, 26 of the 101 *f*_*d*_ blocks overlap with the 9553 HYBRIDCHECK blocks to create a total of 130 discrete overlapping sections. The larger number of blocks identified by HYBRIDCHECK, compared to *f*_*d*_ stat, is likely due to the fact that HYBRIDCHECK may incorrectly identify small regions of incomplete lineage sorting as introgressions.

**Figure 4.**
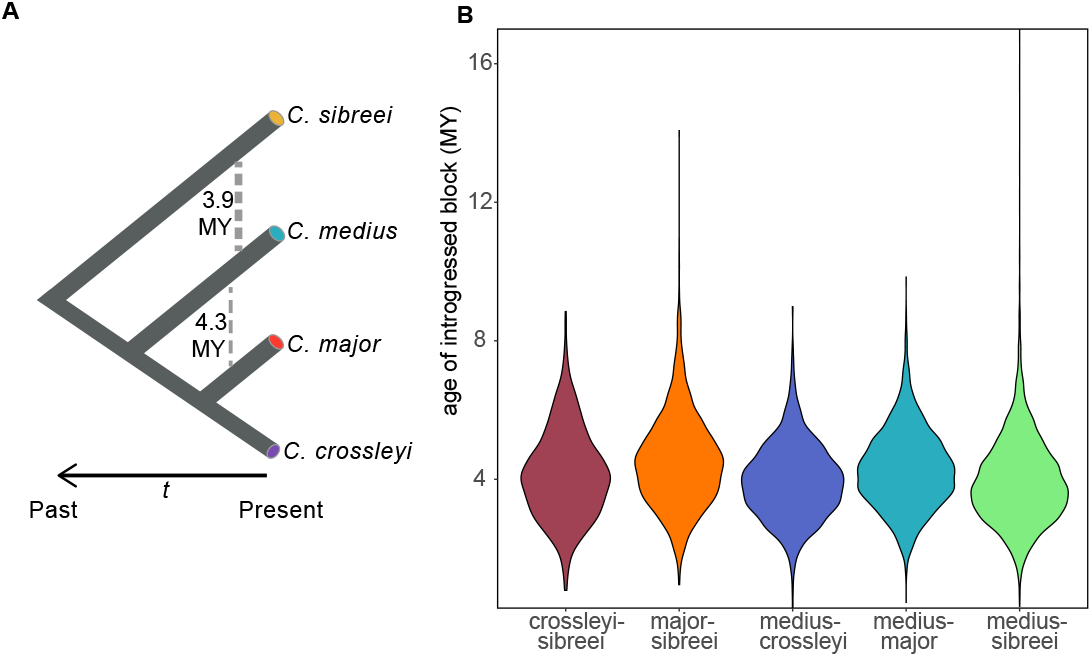
Ancient introgression between *C. sp. cf. medius*, *C. major*, *C. crossleyi* and *C. sibreei*. **A.** Overall relationships within the clade and the approximate mean timing of introgression. Thicker dashed line indicates a larger proportion of admixed regions. **B**. The age of introgressed regions between species pairs.

**Table 1.** Tract sizes and estimated dates of introgression. The mean size of introgressed regions in base pairs is shown above the diagonal and the estimated mean age of introgressed regions in years below the diagonal.

|  | *C. sp. cf. medius* | *C. major* | *C. crossleyi* | *C. sibreei* |
| --- | --- | --- | --- | --- |
| *C. sp. cf. medius* | . | 2,774 | 2,408 | 1,931 |
| *C. major* | 4,280,308 | . | . | 2,240 |
| *C. crossleyi* | 3,941,566 | . | . | 2,116 |
| *C. sibreei* | 3,945,108 | 4,519,688 | 4,135,655 | . |

### Genes within introgressed regions

Parsed RefSeq ID BLAST results were filtered by bit score and evalue, resulting in 4,668 hits, containing 660 genes for *C. med/C. maj* and 4,605 hits, containing 676 genes, for *C. med/C. sib*. However, hibernation genes were not more likely to be in introgressed regions that we would expect by chance for the *C. med/C. maj* test, in fact there was a slight deficit of hibernation genes in introgressed regions, relative to the total number of hibernation genes (χ^2^= 5.55, df = 1, p = 0.018). Nonetheless, introgressed regions for between *C. medius* and *C. major* contained 3% of all hibernation genes (21 unique genes; Supplementary table 3). The *C. med/C. sib* test also did not contain more hibernation genes than expected by chance (Chi-sq = 3.11, df = 1, p = 0.078) and introgressed regions here contained 3.6% of total hibernation genes (25 unique genes; Supp. Table 3). Results do not account for the fact that genes may be non-randomly clustered because annotations are not yet available for the full genome.

GO analyses did detect 30 and 32 GO categories that were significantly over represented in the introgressed regions (p<0.00006, p<0.00009) in the *C. med/C. maj* and *C. med/C. sib* tests, respectively (Supp. Table 4). Among the significantly over-represented GO categories were a large proportion of categories related to transcriptional activity, which is a critical component for inducing the hibernation phenotype (Morin and Storey 2009). Moreover, several genes that appear to have introgressed between *C. medius and C. sibreei* are more directly relevant to hibernation (Fig. 3C). These include genes relating to circadian rhythm, the regulation of feeding (NPY2R; Soscia and Harrington 2005), (HCRTR2; Kilduff and Peyron 2000)), and macrophage lipid homeostasis (ABCA10; Wenzel et al. 2003).

### Historical and contemporary effective population size

To understand how historical demographic events have influenced effective population size (N_e_) through time, we reconstructed the demographic history of each species based on a multiple sequentially Markovian coalescent model (Fig. 5B). From our results, we infer that *Cheirogaleus sp. cf. medius* has seen two expansions in N_e_ in the last two million years, followed by a decline at ~300 kya, with an overall harmonic N_e_ mean of 92,285 individuals. *Cheirogaleus major* shows an initial peak over two million years ago, followed by a decline and then a more moderate N_e_ throughout the last 200 ky, averaging overall N_e_ as 98,005. This trajectory is very similar to that shown by *Cheirogaleus sibreei*, which shows the most stable N_e_ of the *Cheirogaleus* species tested and a relatively high harmonic mean of 103,630. Finally, *C. crossleyi* shows a relatively large increase in N_e_ to almost 200 k; this change occurs at a similar time that we see the decline in *C. sp. cf. medius*. Contemporary heterozygosity is consistent with the MSMC2 results and allows us to infer that the *C. sp. cf. medius* population has not recovered from the decline 300 kya (Fig. 5b). Similarly, the effects of a large increase in *N*_*e*_ seen in *C. crossleyi* are still evident today; the harmonic mean estimate is the highest of the four species, at 108,229 individuals.

**Figure 5.**
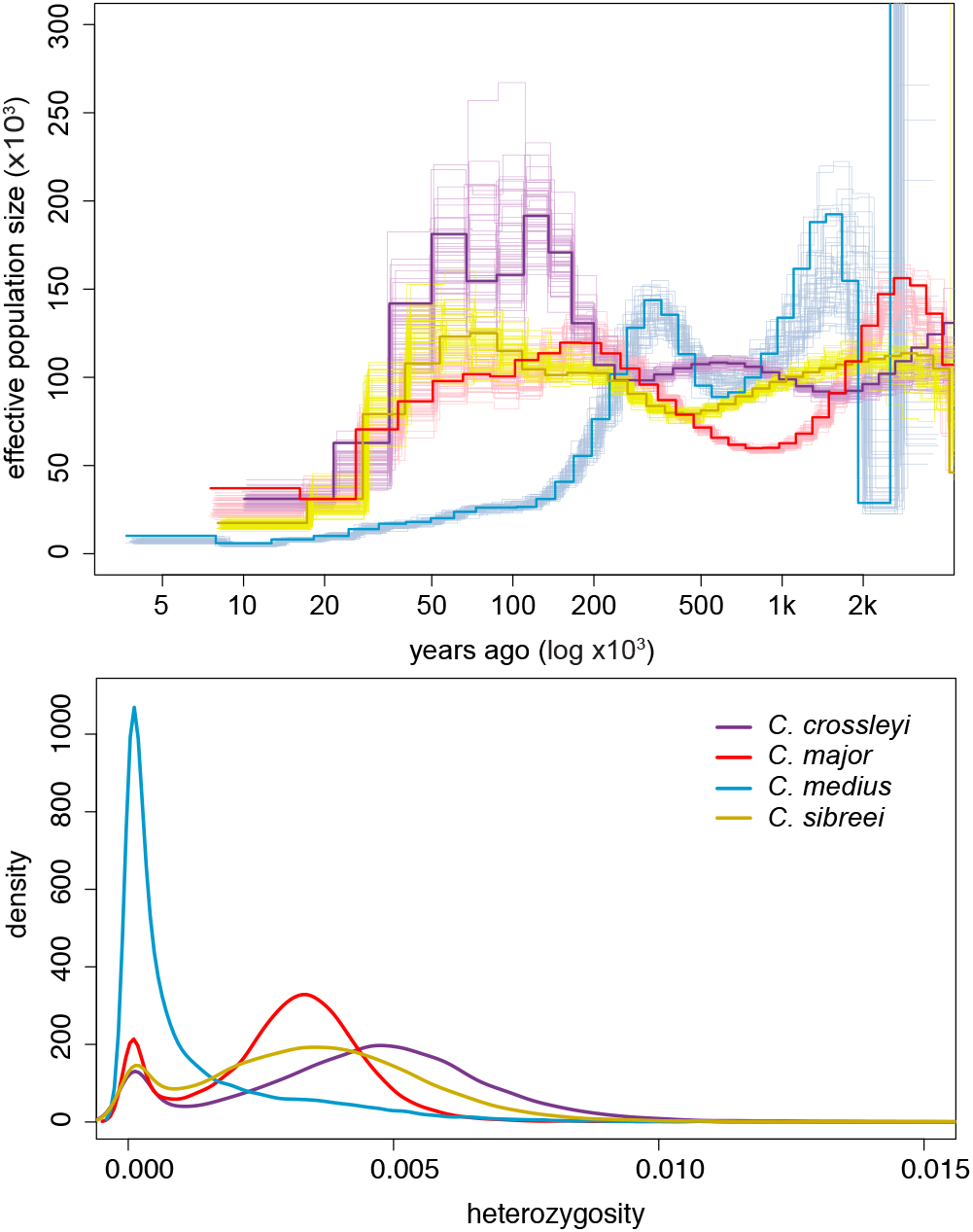
Estimated population sizes and levels of diversity. **A.** Effective population size (N_e_; y-axis) estimated for each species over the last 2 million years (x-axis). The bold line represents the MSMC2 estimate for the original data, and the surrounding lines in the corresponding species colors represent 50 bootstrap replicates. Present time is at the origin and time into the past along the x-axis. **B.** Heterozygosity calculated in 100 kb windows across the genome.

We next quantified heterozygosity for each individual in our data set to test whether long-term population size is correlated with contemporary levels of genetic diversity. We assessed levels of heterozygosity in sliding windows across the genome. The *C. sp. cf. medius* individual included in our study has markedly lower heterozygosity than the other three species (genome-wide average = 0.001 versus 0.003, 0.003, and 0.004; Fig. 5A), while *C. major* and *C. sibreei* both show immediate levels of current heterozygosity (both average 0.003; Fig. 5b). *C. crossleyi* appears to have had a much larger *N*_*e*_ for approximately 150 ky, and this is reflected in it having the highest level of genome-wide heterozygosity (0.004).

## Discussion

Our study not only confirms relationships between Madagascar’s rarely seen nocturnal hibernators, but also identifies ancient introgression that has occurred between species. By utilizing a combination of whole genome sequencing, Markovian coalescent approaches, and phylogenomic analyses of four species of *Cheirogaleus*, we have been able to examine the demographic history of dwarf lemurs in greater resolution than has yet been attempted. We confirmed previously hypothesized phylogenetic relationships among four species within the genus, calculated heterozygosity levels, and assessed historical effective population sizes, thereby using different measures of diversity to assess conservation needs. Surprisingly, these analyses also identified regions of introgression among species that contained several significantly over-represented gene ontology categories.

### Functional implications of ancient admixture for the hibernation phenotype

We identified regions of admixture between *C. sp. cf. medius* and two other species, *C. major* and *C. sibreei*. Contemporary or even relatively recent gene flow between *C. medius* and *C. major* would seem unlikely given the long divergence time (Dos Reis et al. 2018), current morphological differences (including a threefold difference in weight), and the disjunct geographic ranges (dry western forest versus eastern tropical rainforest) of *C. medius* and *C. major* (Goodman and Benstead 2003; IUCN Area of Occupancy distributions shown in Fig. 1A). However, the *C. sp. cf. medius* individual characterized by this study was sampled from Tsihomanaomby, a forest outside of the typical range of the species where it occurs sympatrically with a population of *C. major*. As would be expected given the age of lineage diversification, these introgressed tracts are much smaller than the more recently admixed human genome, where introgressed haplotypes average 50 kb (Sankararaman et al. 2016; Browning et al. 2018). Both the small tract lengths (on average 2.2 kb) and estimates of admixture timing based on sequence divergence indicate that admixture between the species was ancient. Specifically, estimates of the timing of admixture suggests that admixture occurred not long after divergence of the species (Dos Reis et al. 2018).

Building on these estimations of ancient admixture, introgression between *Cheirogaleus medius* and *C. sibreei* may have more biological relevance for understanding their life history. They are the two ‘super hibernators’ in the clade and have similar hibernation profiles relative to the other two species, consistently showing the longest duration of torpor (Blanco et al. 2016, 2018). Both species hibernate for the longest periods of time (up to seven months of the year) and in the more extreme environments: *C. medius* can hibernate under extreme fluctuations in daily ambient temperatures up to 35 °C, whereas *C. sibreei* is a high elevation species that hibernates in the coldest habitats of Madagascar, with an average body temperature of 15 °C (Blanco et al. 2018). They are not sister species, and are estimated to have diverged from the most recent common ancestor 18 Mya (Dos Reis et al. 2018). Hibernation within the clade is hypothesized to be ancestral (Blanco et al. 2018), therefore, the adaptive introgression of genes relating to hibernation is plausible but would have required a strong selective advantage for these regions in the recipient population. Confirmation of adaptive introgression would require additional evidence, such as population-level data showing evidence for a sweep of the introgressed allele (Arnold et al. 2016). The significant GO categories relating to transcriptional activity are particularly interesting because of the transcriptional control mandatory for initiating and maintaining the hibernation phenotype. Entry into torpor requires coordinated controls that suppress and reprioritize all metabolic functions including transcription (Morin and Storey 2009).

We found several genes associated with circadian rhythms to be located in introgressed regions. Gene *HCRTR2* plays a role in the regulation of feeding and sleep/wakefulness (Kilduff and Peyron 2000) and was located in both tests (*C. med/C. maj* and *C. med/C. sib)*. *NPY2R* is a gene that was identified to be upregulated in the fat vs torpor phases of Faherty et al. (2018); *NPY2R* was identified in an introgressed region between *C. med/C. sib*. Circadian clocks play an important role in the timing of mechanisms that regulate hibernation (Hut et al. 2014), as does the circannual clock with its impact on seasonal timing (Helm et al. 2013). The role of the circadian rhythm through the *duration* of hibernation in mammals is unclear, with arctic ground squirrels (*Urocitellus parryii*) showing inhibition of circadian clock function during hibernation (Williams et al. 2017). Neuropeptide Y has been shown to modulate mammalian circadian rhythms (Soscia and Harrington 2005) and there has been interest in its potential for anti-obesity drug development (Yulyaningsih et al. 2011) due to increased expression exerting anorexigenic effects (Yi et al. 2018). Thus, the upregulation of *NPY2R* during the process of hibernation and the highly conserved nature of the Y2 receptor warrants further investigation for its application to human medicine. Also of interest, *ABCA10* was identified in Faherty et al. (2016) to be differentially expressed between ‘active1’ and ‘torpor’ states (‘active1’ represents the accumulation of weight during the summer months). This gene is shown to be present in introgressed regions in the *C. med/C. sib* test. *ABCA10* plays a role in macrophage lipid homeostasis (Wenzel et al. 2003). Body fat reserves are essential to enter hibernation and are then metabolised and depleted throughout the season (Blanco et al. 2018). As longer hibernators, *C. medius* and *C. sibreei* would be expected to need greater control over fat metabolism.

### Climatic impacts on population size through time and prospects for the future

Madagascar’s history of climatic variation has shaped the fauna of the island (Dewar and Richard 2007). Fluctuation of climate such as lower temperatures at the Eocene-Oligocene transition (Dupont-Nivet et al. 2007), followed by warming climate and low precipitation of the Miocene, caused dramatic contractions and expansions of vegetation (Hannah et al. 2008). Climatic variation has been incorporated into models that try to explain the micro-endemism and historical patterns of dispersal and vicariance across the island (Wilmé et al. 2014). Martin (1972) posited a model that considered larger rivers as geographical barriers to gene flow, suggesting that climatically and physically defined zones can operate as agents for geographical isolation and speciation, such as climatic pulsing. Wilmé et al. (2006) followed a similar theme looking at watersheds in the context of Quaternary climatic and vegetation shifts, resulting in patterns of diversity and microendemism. Craul et al. (2007) tested the influence of inter-river-systems on simultaneously isolating populations and providing retreat zones.

The changes in N_e_ seen in Figure 4B, fit with Wilmé’s hypothesis of climatic pulsing. During periods of glaciation, when the climate was cooler and drier, high altitude species were likely buffered from changing conditions, whereas low altitude species likely experienced declines associated with aridification and greater levels of habitat isolation, and may have expanded during glacial minima, when climatic conditions were warmer and more humid. The finding of a more stable N_e_ through time for *C. sibreei*, the high plateau specialist, supports the theory that populations at high altitudes avoided the expansions and contractions occurring at lower elevations. Results for species typically found at lower elevations show more variation e.g. a decrease in *C. sp. cf. medius* N_e_ occurs simultaneously with an increase in *C. crossleyi* N_e_. The inference of demographic history is strongly influenced by population structure and changes in connectivity (Hawks 2017; Song et al. 2017), and so we are careful to discuss overall trends and not specifics. Results show an overall decline for all species in the last 50 k years. Declines in more recent times are echoed in great apes, including modern humans (Prado-martinez et al. 2013). While extremes at either end of the time scale in MSMC2 should be taken with caution (Hawks 2017), it appears that certain dwarf lemur populations have been declining for as long as 300 ky. Furthermore, these declines are well supported by the bootstrap values, particularly in the case of *C. sp. cf. medius*.

Our results indicate that population declines of dwarf lemurs occurred long before the arrival of humans, introducing the possibility of long-term low N_e_. As these are all species of conservation concern, we assessed genomic levels of heterozygosity to derive an estimate of genetic diversity. *C. sp. cf. medius* had the lowest overall heterozygosity of the samples here examined, and these values are indicative of a small and isolated population at risk of inbreeding depression (Rogers and Slatkin 2017). The *C. sp. cf. medius* individual examined here was collected from Tsihomanaomby in the north-east of Madagascar. This is a sub-humid rainforest site (~7 km^2^) not typical to the *C. medius* habitat preference given that the species is normally associated with the dry forests of the west (Groves 2000; Hapke et al. 2005; Groeneveld et al. 2010, 2011). Interestingly, our estimates of historic effective population size trajectory suggest that the N_e_ of *C. sp. cf. medius* has long been much lower than that of other dwarf lemurs (Fig. 4B). It is therefore possible that the decline in *C. sp. cf. medius* N_e_ seen ~300 kya is the time that this population became isolated from other *C. medius* populations, or a consequence of a bottleneck associated with their colonisation of Tsihomanaomby. It is plausible that the Tsihomanaomby population was founded by individuals from the north west (from which it now appears separated; Fig. 1B), and that founder effects are still impacting the present-day population, indicated the by low heterozygosity in our *C. sp. cf. medius* individual (Fig. 4A).

### Summary

Our study illustrates the power of selective sampling and genomic analysis for identifying populations/species in need of conservation, given that analyses of whole genomes allows us to identify historical events detectable in the genomes of extant individuals. We conclude from our analyses that the *Cheirogaleus sp. cf. medius* population warrants further investigation, both taxonomically, given its disjunct distribution, and as a species of conservation concern. All four taxa show temporal changes in ancestral N_e_ over the last two million years with an overall decrease in N_e_ in recent years (50 k years). While species may recover from some fluctuation in population size is recoverable, large crashes leave long term traces in the genome that can be detrimental to long term survival of populations (Palkopoulou et al. 2015). We demonstrate that for *Cheirogaleus*, such events can be potentially disadvantageous, such as low heterozygosity from population declines, as seen in *C. sp. cf. medius*. However, the significant enrichment of several categories of genes in introgressed regions of the genome, shown by the GO analysis, demonstrates for the first time a potential source of novel variation in *Cheirogaleus*. Hybridization is a pervasive evolutionary force and can drive adaptive differentiation between populations (Runemark et al. 2018). We show that ancient admixture may have been a possible mechanism for adaptive phenotypic evolution in these species of concern.

## Acknowledgements

We thank Stephen D. Nash for the use of his *Cheirogaleus* illustrations. We thank Professor Manfred Grabherr and Jessica Rick for advice on analyses, and Michelle Larsen for her time in the lab. We thank Rikki Gumbs and three anonymous reviewers for comments on the manuscript. We thank the Malagasy authorities for permission to conduct this research. This study was funded by a National Science Foundation Grant DEB-1354610 and Duke University startup funds to ADY. RCW and ADY also received a Matching Funds Award from Dovetail Genomics for the reference genome. ADY gratefully acknowledges support from the John Simon Guggenheim Foundation and the Alexander von Humboldt Foundation during the writing phase of this project. We are grateful for the support of Duke Research Computing and the Duke Data Commons (NIH 1S10OD018164-01). This is Duke Lemur Center publication no. XXX.

## Data Archiving

Genomic sequences can be found under NCBI BioProject accession PRJNA523575. BioSample accessions are listed in Supplementary Table 1.

## Supplementary Material

**Table S1.** Sequencing information for all samples sequenced

| Species | Sample # | BioProject accession | BioSample accession | Sample Quality (bp) | Library type | Lanes | Pooled | Index sequence | Index name | Insert size | Passed Filter Yield (bp) |
| --- | --- | --- | --- | --- | --- | --- | --- | --- | --- | --- | --- |
| *Cheirogaleus medius* | 106355 | PRJNA523575 | SAMN11334321 | 150,000 | shotgun | 2 |  | GGCTACA | TruSeq #11 | 350 | 208,330,574,982 |
|  |  |  |  |  | Chicago | 2 | pool 1 | CCTAGGT | 3 | 654 | 76,809,567,130 |
|  |  |  |  |  |  |  |  | GGATCAA | 4 | 651 | 68,643,708,496 |
|  |  |  |  |  |  |  |  | TTGGATC | 14 | 649 | 79,318,091,474 |
| *C. sp. cf. medius* | 106295 | PRJNA523575 | SAMN11334317 | 19,183 | DNA-Seq | 4 | pool 2 | CGGCTATG |  | 300 | 108,952,269,408 |
| *C. major* | 106242 | PRJNA523575 | SAMN11334318 | >60,000 |  |  |  | TCCGCGAA |  |  | 123,181,913,260 |
| *C. crossleyi* | 106209 | PRJNA523575 | SAMN11334319 | 56,688 |  |  |  | TCTCGCGC |  |  | 98,250,428,892 |
| *C. sibreei* | 106186 | PRJNA523575 | SAMN11334320 | 21,550 |  |  |  | AGCGATAG |  |  | 117,625,961,276 |

**Table S2.**
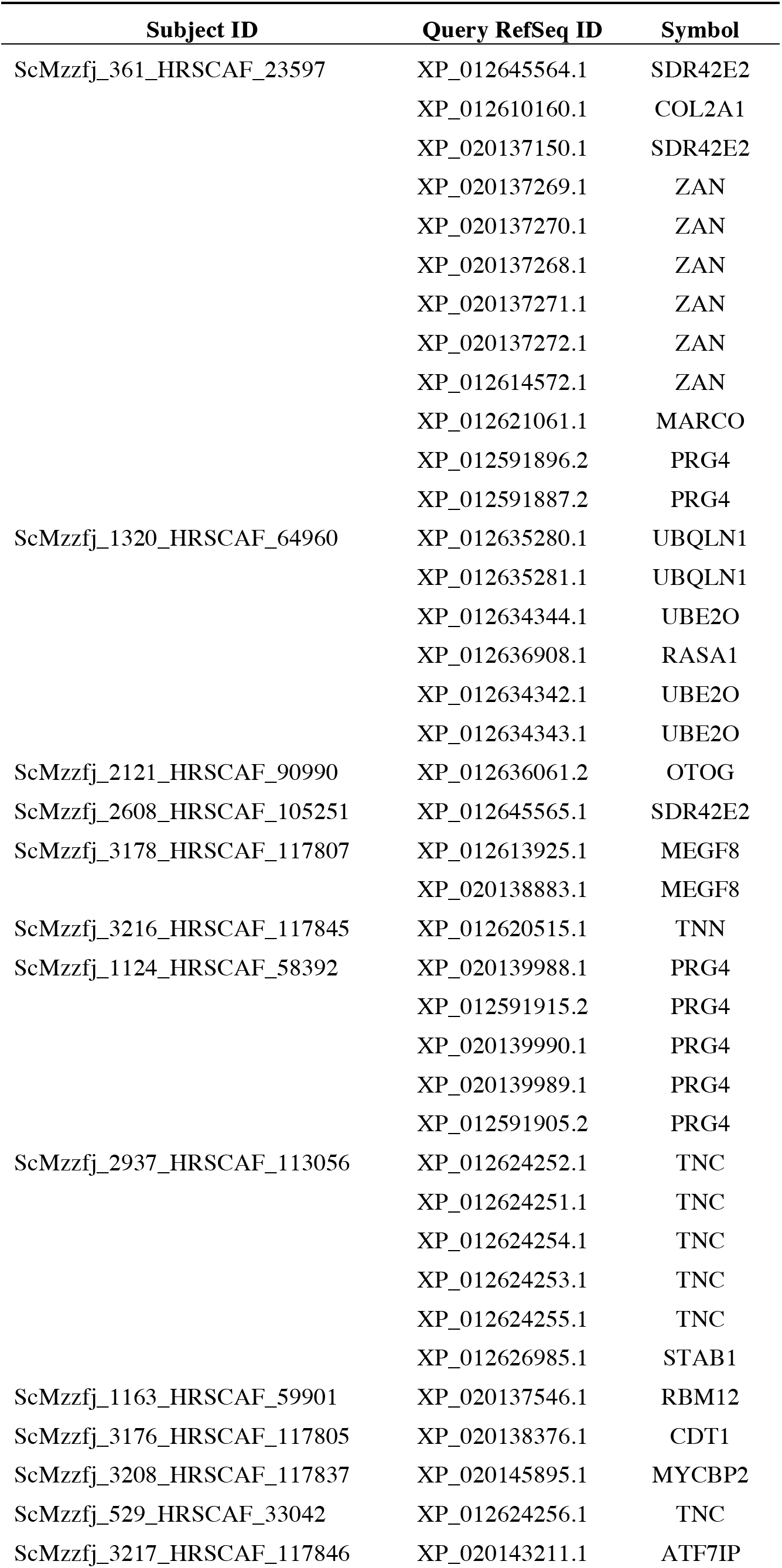
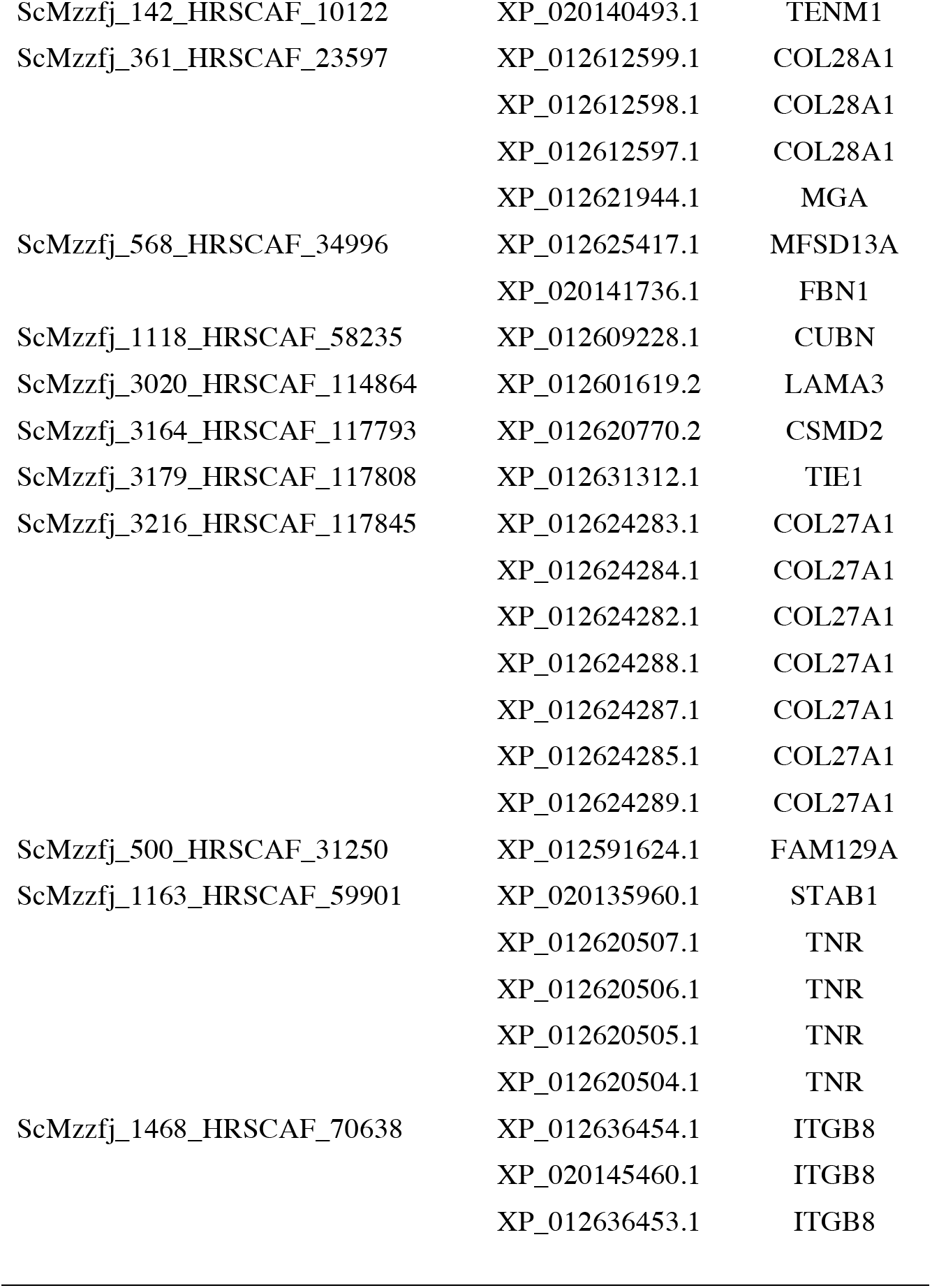
Table shows regions that have associated gene names identified in SAGUARO topologies that do not agree with the previously accepted taxonomic relationship within *Cheirogaleus*. Regions without associated gene names have been excluded here.

**Table S3.**
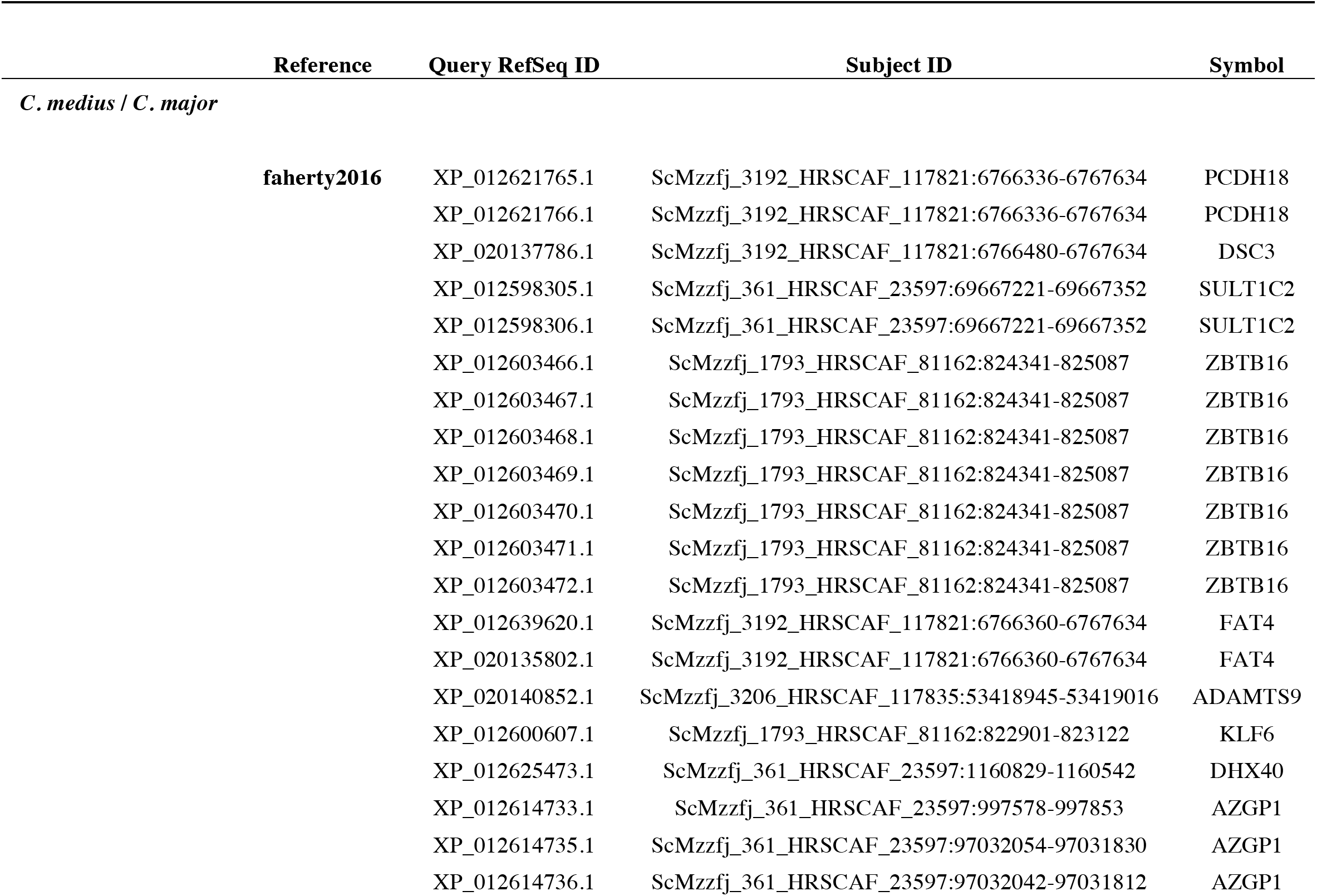
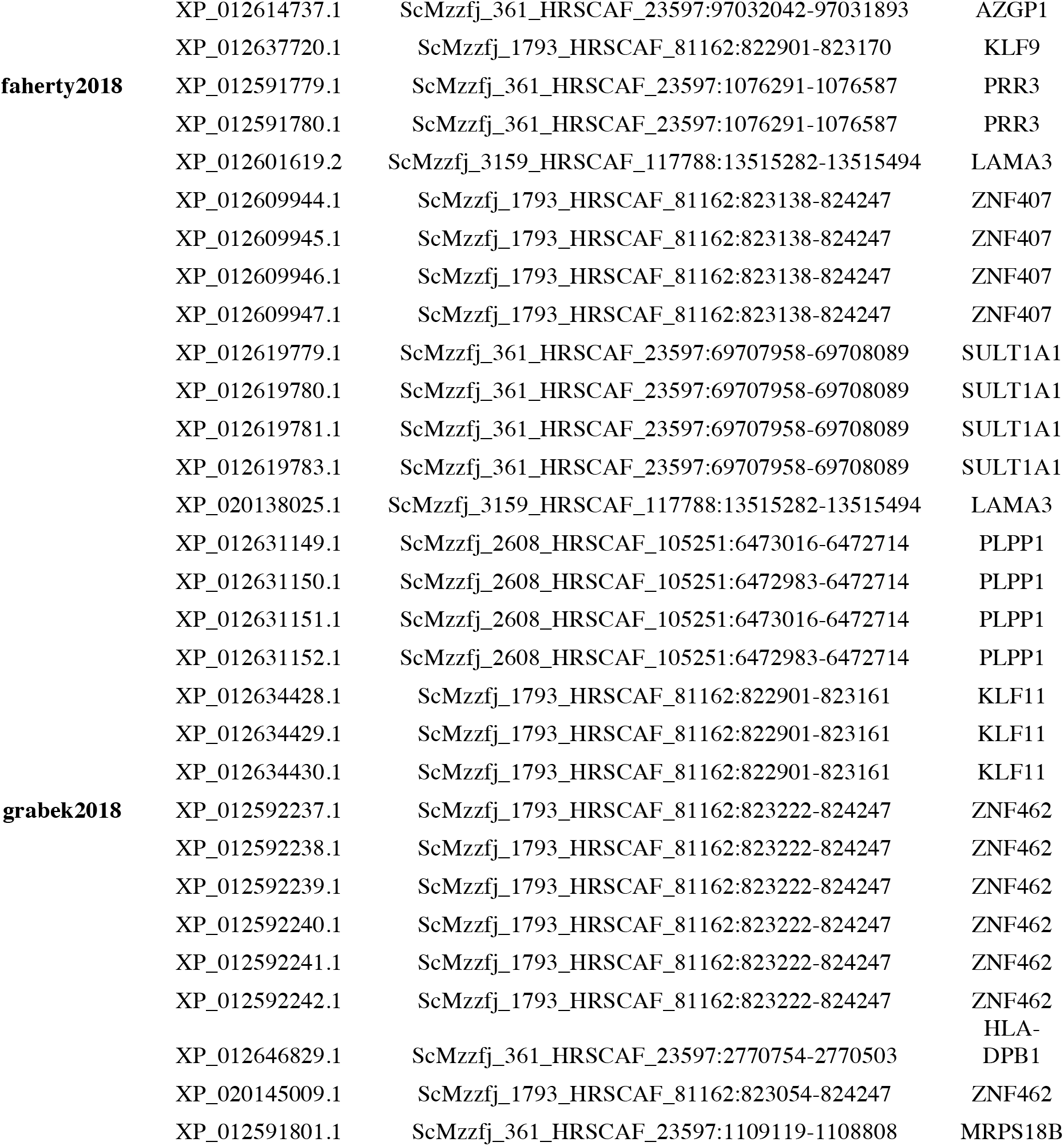
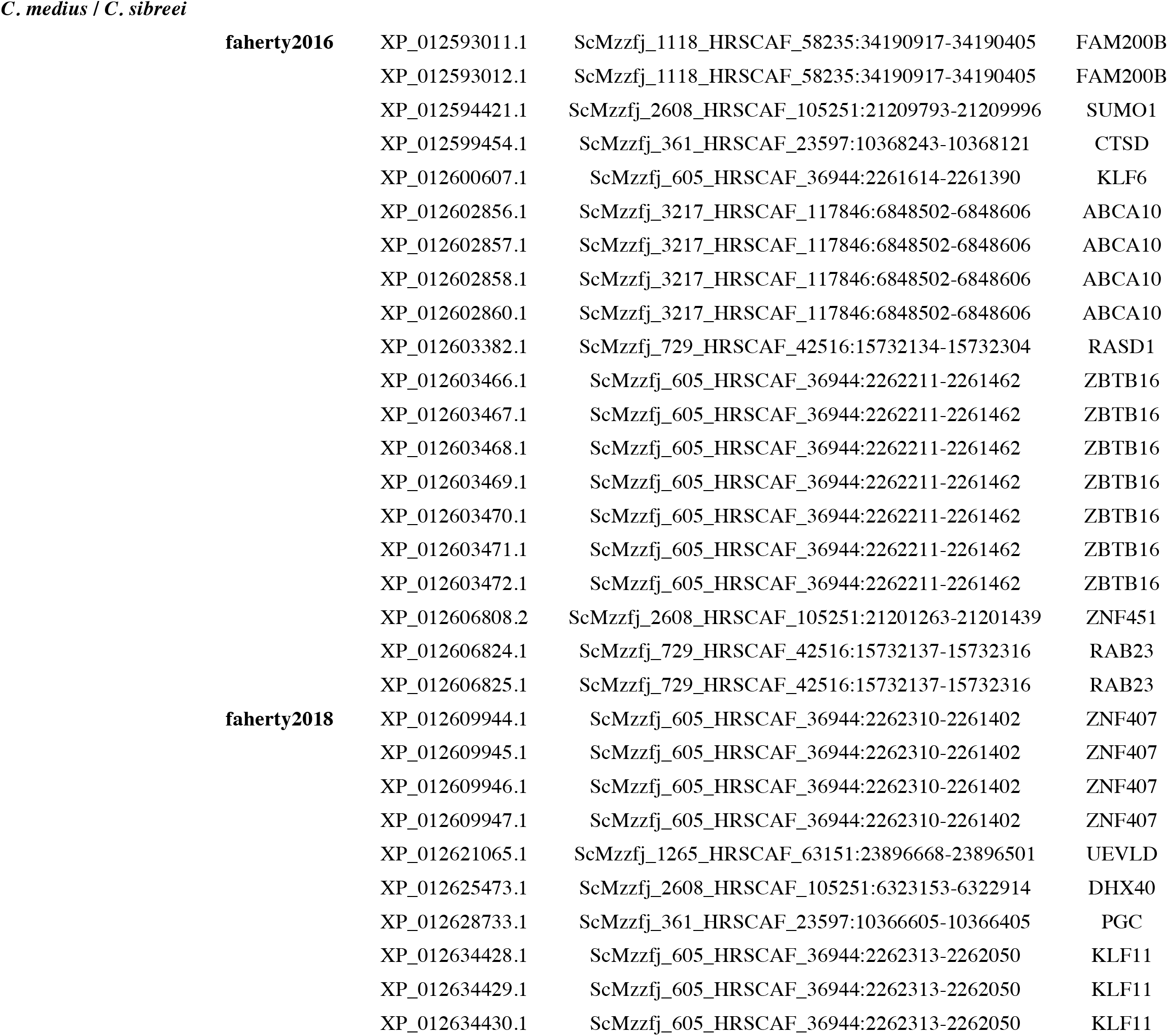
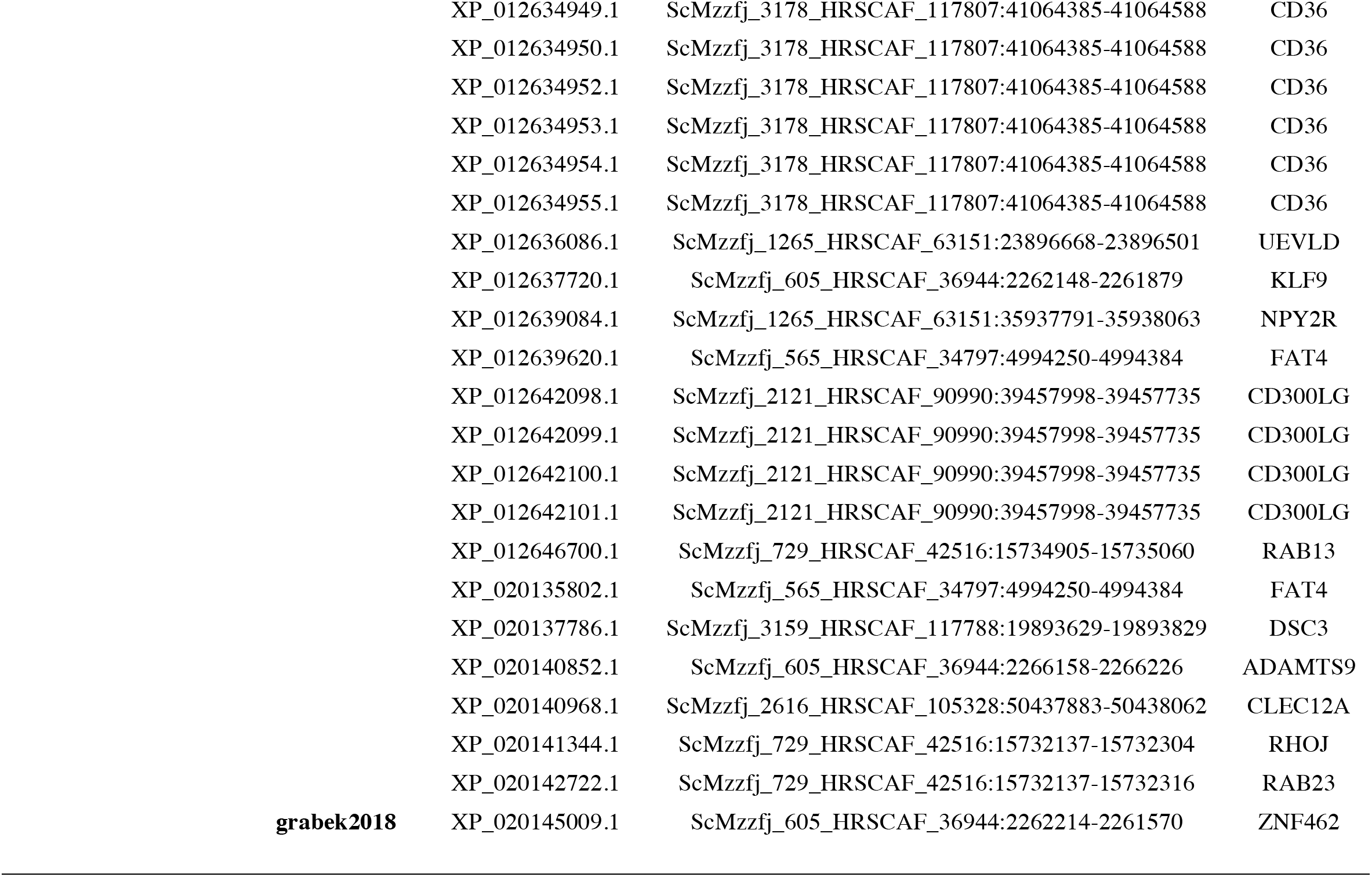
Regions of the genome shown to have elevated *f*_*d*_ values that also contained genes associated with hibernation (Faherty et al. 2016, 2018; Grabek et al. 2017). The reference identifying the genes involvement in hibernation is shown. Query ID refers to the *Microcebus murinus* RefSeq ID, Subject ID refers to the scaffold and region of the *Cheirogaleus* genome.

**Table S4.** A summary of gene ontology results for introgressed regions of the genome for tests between *Cheirogaleus sp. cf. medius /C. major*, and C. *sp. cf. medius*/*C. sibreei*. All results are significantly over enriched for both tests respectively (p<0.00006, p<0.00009).

| FDR |  |  |
| --- | --- | --- |
| *C. med/C. maj* | *C. med/C. sib* | GO description |
| 7.30E-20 | 2.57E-20 | negative regulation of transcription from RNA polymerase II promoter |
| 1.70E-13 | 2.41E-13 | transcriptional repressor activity, RNA polymerase II core promoter proximal region sequence-specific binding |
| 1.88E-08 | 2.53E-09 | transcriptional repressor activity, RNA polymerase II transcription regulatory region sequence-specific binding |
| 8.23E-05 | 4.34E-07 | positive regulation of transcription from RNA polymerase II promoter |
| 7.19E-05 | 9.62E-06 | negative regulation of transcription, DNA-templated |
| 0.00134417 | 6.72E-05 | transcription factor activity, sequence-specific DNA binding |
| 4.44E-05 | 0.00020978 | transcriptional activator activity, RNA polymerase II transcription regulatory region sequence-specific binding |
|  | 0.000708435 | transcription regulatory region DNA binding |
| 8.25E-05 | 0.001018928 | positive regulation of transcription, DNA-templated |
|  | 0.002931254 | transcriptional repressor activity, RNA polymerase II distal enhancer sequence-specific binding |
| 0.006679222 | 0.009879882 | regulation of transcription, DNA-templated |
| 0.02581555 | 0.038651284 | transcriptional activator activity, RNA polymerase II core promoter proximal region sequence-specific binding |
|  | 0.040293509 | transcriptional repressor complex |
| 0.000135577 |  | transcription regulatory region DNA binding |

## References

Abascal, F., A. Corvelo, F. Cruz, J. L. Villanueva-Cañas, A. Vlasova, M. Marcet-Houben, B. Martínez-Cruz, J. Y. Cheng, P. Prieto, V. Quesada, J. Quilez, G. Li, F. García, M. Rubio-Camarillo, L. Frias, P. Ribeca, S. Capella-Gutiérrez, J. M. Rodríguez, F. Câmara, E. Lowy, L. Cozzuto, I. Erb, M. L. Tress, J. L. Rodriguez-Ales, J. Ruiz-Orera, F. Reverter, M. Casas-Marce, L. Soriano, J. R. Arango, S. Derdak, B. Galán, J. Blanc, M. Gut, B. Lorente-Galdos, M. Andrés-Nieto, C. López-Otín, A. Valencia, I. Gut, J. García, R. Guigó, W. J. Murphy, A. Ruiz-Herrera, T. Marques-Bonet, G. Roma, C. Notredame, T. Mailund, M. M. Albà, T. Gabaldón, T. Alioto, and J. A. Godoy. 2016. Extreme genomic erosion after recurrent demographic bottlenecks in the highly endangered Iberian lynx. Genome Biol. 17. Genome Biology.

Albert-Daviaud, A., S. Perillo, and W. Stuppy. 2018. Seed dispersal syndromes in the Madagascan flora: The unusual importance of primates. Oryx 52:418–426.

Altschul, S. F., W. Gish, W. Miller, E. W. Myers, and D. J. Lipman. 1990. Basic local alignment search tool. J. Mol. Biol. 215:403–410.

Árnason, Ú., F. Lammers, V. Kumar, M. A. Nilsson, and A. Janke. 2018. Whole-genome sequencing of the blue whale and other rorquals finds signatures for introgressive gene flow. Sci. Adv. 4.

Arnold, B. J., B. Lahner, J. M. Dacosta, C. M. Weisman, J. D. Hollister, and D. E. Salt. 2016. Borrowed alleles and convergence in serpentine adaptation. PNAS 113:1–6.

Blanco, M. B., K. H. Dausmann, S. L. Faherty, P. Klopfer, A. D. Krystal, R. Schopler, and A. D. Yoder. 2016. Hibernation in a primate: Does sleep occur? R. Soc. Open Sci. 3.

Blanco, M. B., K. H. Dausmann, S. L. Faherty, and A. D. Yoder. 2018. Tropical heterothermy is “cool”: The expression of daily torpor and hibernation in primates. Evol. Anthropol. 147–161.

Blanco, M. B., L. R. Godfrey, M. Rakotondratsima, V. Rahalinarivo, K. E. Samonds, J. L. Raharison, and M. T. Irwin. 2009. Discovery of sympatric dwarf lemur species in the high-altitude rain forest of Tsinjoarivo, Eastern Madagascar: Implications for biogeography and conservation. Folia Primatol. 80:1–17.

Bolger, A. M., M. Lohse, and B. Usadel. 2014. Trimmomatic: A flexible trimmer for Illumina sequence data. Bioinformatics 30:2114–2120.

Broad Institute. 2018. Picard.

Browning, S. R., B. L. Browning, Y. Zhou, S. Tucci, J. M. Akey, S. R. Browning, B. L. Browning, Y. Zhou, S. Tucci, and J. M. Akey. 2018. Analysis of Human Sequence Data Reveals Two Pulses of Archaic Denisovan Admixture Article Analysis of Human Sequence Data Reveals Two Pulses of Archaic Denisovan Admixture. Cell 173:53–61.e9. Elsevier.

Burns, S. J., L. R. Godfrey, P. Faina, D. McGee, B. Hardt, L. Ranivoharimanana, and J. Randrianasy. 2016. Rapid human-induced landscape transformation in Madagascar at the end of the first millennium of the Common Era. Quat. Sci. Rev. 134:92–99. Elsevier Ltd.

Ceballos, G., and P. Ehrlich. 2018. The misunderstood sixth mass extinction. Science (80-.). 360:1080–1081.

Ceballos, G., P. R. Ehrlich, A. D. Barnosky, A. García, R. M. Pringle, and T. M. Palmer. 2015. Accelerated modern human – induced species losses: entering the sixth mass extinction. Sci. Adv. 1:1–5.

Chen, N., E. J. Cosgrove, R. Bowman, J. W. Fitzpatrick, A. G. Clark, N. Chen, E. J. Cosgrove, R. Bowman, J. W. Fitzpatrick, and A. G. Clark. 2016. Genomic Consequences of Population Decline in the Report Genomic Consequences of Population Decline in the Endangered Florida Scrub-Jay. Curr. Biol. 26:1–6. Elsevier Ltd.

Craul, M., E. Zimmermann, S. Rasoloharijaona, B. Randrianambinina, and U. Radespiel. 2007. Unexpected species diversity of Malagasy primates (Lepilemur spp.) in the same biogeographical zone: a morphological and molecular approach with the description of two new species. 15:1–15.

Depristo, M. A., E. Banks, R. Poplin, K. V. Garimella, J. R. Maguire, C. Hartl, A. A. Philippakis, G. Del Angel, M. A. Rivas, M. Hanna, A. McKenna, T. J. Fennell, A. M. Kernytsky, A. Y. Sivachenko, K. Cibulskis, S. B. Gabriel, D. Altshuler, and M. J. Daly. 2011. A framework for variation discovery and genotyping using next-generation DNA sequencing data. Nat. Genet. 43:491–501.

Dewar, R. E., and A. F. Richard. 2007. Evolution in the hypervariable environment of Madagascar Madagascar. PNAS 104:13723–13727.

Dirzo, R., H. S. Young, M. Galetti, G. Ceballos, N. J. B. Isaac, and B. Collen. 2014. Defaunation in the Anthropocene. Science (80-.). 345:401–406.

Dos Reis, M., G. F. Gunnell, J. Barba-Montoya, A. Wilkins, Z. Yang, and A. D. Yoder. 2018. Using phylogenomic data to explore the effects of relaxed clocks and calibration strategies on divergence time estimation: Primates as a test case. Syst. Biol. 67:594–615.

Dupont-Nivet, G., W. Krijgsman, C. G. Langereis, H. A. Abels, S. Dai, and X. Fang. 2007. Tibetan plateau aridification linked to global cooling at the Eocene-Oligocene transition. Nature 445:635–638.

Durand, E. Y., N. Patterson, D. Reich, and M. Slatkin. 2011. Testing for ancient admixture between closely related populations. Mol. Biol. Evol. 28:2239–2252.

Durinck, S., Y. Moreau, A. Kasprzyk, S. Davis, B. De Moor, A. Brazma, and W. Huber. 2005. BioMart and Bioconductor: a powerful link between biological databases and microarray data analysis. Bioinformatics 3439–3440.

Durinck, S., P. Spellman, E. Birney, and W. Huber. 2009. Mapping identifiers for the integration of genomic datasets with the R/Bioconductor package biomaRt. Nat. Protoc. 1184–1191.

Estrada, A., P. A. Garber, A. B. Rylands, C. Roos, E. Fernandez-duque, A. Di Fiore, K. A. Nekaris, V. Nijman, E. W. Heymann, J. E. Lambert, F. Rovero, C. Barelli, and L. Baoguo. 2017. Impending extinction crisis of the world’s primates: Why primates matter. Sci. Adv. 3:e1600946.

Faherty, S. L., J. L. Villanueva-Cañas, M. B. Blanco, M. M. Albà, and A. D. Yoder. 2018. Transcriptomics in the wild: Hibernation physiology in free-ranging dwarf lemurs. Mol. Ecol. 27:709–722.

Faherty, S. L., J. L. Villanueva-Cañas, P. H. Klopfer, M. M. Albà, and A. D. Yoder. 2016. Gene expression profiling in the hibernating primate, *Cheirogaleus medius*. Genome Biol. Evol. evw163.

Figueiró, H. V., G. Li, F. J. Trindade, J. Assis, F. Pais, G. Fernandes, S. H. D. Santos, G. M. Hughes, A. Komissarov, A. Antunes, C. S. Trinca, M. R. Rodrigues, T. Linderoth, K. Bi, L. Silveira, F. C. C. Azevedo, D. Kantek, E. Ramalho, R. A. Brassaloti, P. M. S. Villela, A. L. V. Nunes, R. H. F. Teixeira, R. G. Morato, D. Loska, P. Saragüeta, T. Gabaldón, E. C. Teeling, S. J. O’Brien, R. Nielsen, L. L. Coutinho, G. Oliveira, W. J. Murphy, and E. Eizirik. 2017. Genome-wide signatures of complex introgression and adaptive evolution in the big cats. Sci. Adv. 3:e1700299.

Frasier, C. L., R. Lei, A. T. Mclain, J. M. Taylor, C. A. Bailey, A. L. Ginter, S. D. Nash, R. Randriamampionona, C. P. Groves, R. A. Mittermeier, and E. E. L. Jr. 2016. A New Species of Dwarf Lemur (Cheirogaleidae: Cheirogaleus medius Group) from the Ankarana and AndrafiamenaAandavakoera Massifs, Madagascar. Primate Conserv. 2016:59–72.

Goodman, S., and J. Benstead. 2003. The natural history of Madagascar. The University of Chicago Press.

Goodman, S., and J. Bernstead. 2007. The Natural History of Madagascar. Illustrate. University of Chicago Press.

Grabek, K. R. K., T. T. F. Cooke, L. E. Epperson, K. S.- BioRxiv, U. 2017, K. K. Spees, G. F. Cabral, S. C. Sutton, D. K. Merriman, S. L. Martin, and C. D. Bustamante. 2017. Genetic architecture drives seasonal onset of hibernation in the 13-lined ground squirrel. bioRxiv 1–36.

Green, R. E., J. Krause, A. W. Briggs, T. Maricic, U. Stenzel, M. Kircher, N. Patterson, H. Li, W. Zhai, M. H. Fritz, N. F. Hansen, E. Y. Durand, A. Malaspinas, J. D. Jensen, T. Marques-bonet, C. Alkan, K. Prüfer, M. Meyer, H. A. Burbano, J. M. Good, R. Schultz, A. Aximu-petri, A. Butthof, B. Höber, B. Höffner, M. Siegemund, A. Weihmann, C. Nusbaum, E. S. Lander, C. Russ, N. Novod, J. Affourtit, M. Egholm, J. Fortea, A. Rosas, R. W. Schmitz, P. L. F. Johnson, E. E. Eichler, D. Falush, E. Birney, J. C. Mullikin, M. Slatkin, R. Nielsen, J. Kelso, M. Lachmann, D. Reich, and S. Pääbo. 2010a. A Draft Sequence of the Neandertal Genome. Science (80-.). 328:710–723.

Green, R., J. Krause, A. W. Briggs, T. Maricic, U. Stenzel, M. Kircher, N. Patterson, H. Li, W. Zhai, M. H.-Y. Fritz, N. F. Hansen, E. Y. Durand, A.-S. Malaspinas, J. D. Jensen, and T. Marques-Bonet. 2010b. A draft sequence of the neadertal genome. Science (80-.). 328:710–722.

Groeneveld, L. F., M. B. Blanco, J. L. Raharison, V. Rahalinarivo, R. M. Rasoloarison, P. M. Kappeler, L. R. Godfrey, and M. T. Irwin. 2010. MtDNA and nDNA corroborate existence of sympatric dwarf lemur species at Tsinjoarivo, eastern Madagascar. Mol. Phylogenet. Evol. 55:833–845. Elsevier Inc.

Groeneveld, L. F., R. M. Rasoloarison, and P. M. Kappeler. 2011. Morphometrics confirm taxonomic deflation in dwarf lemurs (Primates: Cheirogaleidae), as suggested by genetics. Zool. J. Linn. Soc. 161:229–244.

Groeneveld, L., D. Weisrock, R. Rasoloarison, A. Yoder, and P. Kappeler. 2009. Species delimitation in lemurs: multiple genetic loci reveal low levels of species diversity in the genus Cheirogaleus. BMC Evol. Biol. 9:1–16.

Groves, C. P. 2000. The genus Cheirogaleus: Unrecognized biodiversity in dwarf lemurs. Int. J. Primatol. 21:943–962.

Haddad, N. M., L. A. Brudvig, J. Clobert, K. F. Davies, A. Gonzalez, R. D. Holt, T. E. Lovejoy, J. O. Sexton, M. P. Austin, C. D. Collins, W. M. Cook, E. I. Damschen, R. M. Ewers, B. L. Foster, C. N. Jenkins, A. J. King, W. F. Laurance, D. J. Levey, C. R. Margules, B. A. Melbourne, A. O. Nicholls, J. L. Orrock, D. Song, and J. R. Townshend. 2015. Habitat fragmentation and its lasting impact on Earth‘s ecosystems. Appl. Ecol. 1–9.

Hajdinjak, M., Q. Fu, A. Hübner, M. Petr, F. Mafessoni, S. Grote, P. Skoglund, V. Narasimham, H. Rougier, I. Crevecoeur, P. Semal, M. Soressi, S. Talamo, J. J. Hublin, I. Gušić, Z. Kućan, P. Rudan, L. V. Golovanova, V. B. Doronichev, C. Posth, J. Krause, P. Korlević, S. Nagel, B. Nickel, M. Slatkin, N. Patterson, D. Reich, K. Prüfer, M. Meyer, S. Pääbo, and J. Kelso. 2018. Reconstructing the genetic history of late Neanderthals. Nature 555:652–656.

Hannah, L., R. Dave, P. Lowry, S. Andelmann, M. Andrianarisata, L. Andriamaro, A. Cameron, R. Hijmans, C. Kreman, M. J, H. Randrianasolo, S. Andriambololonera, A. Razafimpahanan, H. Randriamahazo, J. Randrianarisoa, P. Razafinjatov, C. Raxworthy, G. Schatz, M. Tadross, and L. Wilmé. 2008. Climate change adaptation for conservation in Madagascar. Biol. Lett. 4:590–594.

Hapke, A., J. Fietz, S. D. Nash, D. Rakotondravony, B. Rakotosamimanana, J. B. Ramanamanjato, G. F. N. Randria, and H. Zischler. 2005. Biogeography of dwarf lemurs: Genetic evidence for unexpected patterns in southeastern Madagascar. Int. J. Primatol. 26:873–901.

Harris, K., and R. Nielsen. 2016. The Genetic Cost of introgression. Genetics http://dx.doi.org/10.1534/genetics.116.186890.

Harrison, R. G., and E. L. Larson. 2014. Hybridization, introgression, and the nature of species boundaries. J. Hered. 105:795–809.

Hawks, J. 2017. Introgression makes waves in inferred histories of effective population size. 89:1–38.

Helm, B., R. Ben-Shlomo, M. J. Sheriff, R. A. Hut, R. Foster, B. M. Barnes, and D. Dominoni. 2013. Annual rhythms that underlie phenology: Biological time-keeping meets environmental change. Proc. R. Soc. B Biol. Sci. 280.

Hoang, D. T., O. Chernomor, A. Von Haeseler, B. Q. Minh, and L. S. Vinh. 2018. UFBoot2: Improving the ultrafast bootstrap approximation. Mol. Biol. Evol. 35:518–522.

Hut, R. A., H. Dardente, and S. J. Riede. 2014. Seasonal timing: How does a hibernator know when to stop hibernating? Curr. Biol. 24:R602–R605. Elsevier.

IUCN. 2019. The IUCN Red List of Threatened Species.

Kalyaanamoorthy, S., B. Q. Minh, T. K. F. Wong, A. Von Haeseler, and L. S. Jermiin. 2017. ModelFinder: Fast model selection for accurate phylogenetic estimates. Nat. Methods 14:587–589.

Katoh, K., K. Misawa, K. Kuma, and T. Miyata. 2002. MAFFT: a novel method for rapid multiple sequence alignment based on fast Fourier transform. Nucleic Acids Res. 30:3059–3066.

Kilduff, T. S., and C. Peyron. 2000. The hypocretin/orexin ligand-receptor system: Implications for sleep and sleep disorders. Trends Neurosci. 23:359–365.

Kimura, M. 1968. Evolutionary rate at the molecular level. Nature 217:624–626.

Korneliussen, T. S., A. Albrechtsen, and R. Nielsen. 2014. ANGSD: Analysis of Next Generation Sequencing Data. BMC Bioinformatics 15:1–13.

Larsen, P. A., R. A. Harris, Y. Liu, S. C. Murali, C. R. Campbell, A. D. Brown, B. A. Sullivan, J. Shelton, S. J. Brown, M. Raveendran, O. Dudchenko, I. Machol, N. C. Durand, M. S. Shamim, E. L. Aiden, D. M. Muzny, R. A. Gibbs, A. D. Yoder, J. Rogers, and K. C. Worley. 2017. Hybrid de novo genome assembly and centromere characterization of the gray mouse lemur (Microcebus murinus). 1–17. BMC Biology.

Lei, R., C. L. Frasier, A. T. McLain, J. M. Taylor, C. A. Bailey, S. E. Engberg, A. L. Ginter, R. Randriamampionona, C. P. Groves, R. A. Mittermeier, and E. E. L. Jr. 2014. Revision of Madagascar’s Dwarf Lemurs (Cheirogaleidae: Cheirogaleus): Designation of Species, Candidate Species Status and Geographic Boundaries Based on Molecular and Morphological Data. Primate Conserv. 28:9–35.

Lei, R., A. T. McLain, C. L. Frasier, J. M. Taylor, C. A. Bailey, S. E. Engberg, A. L. Ginter, S. D. Nash, R. Randriamampionona, C. P. Groves, R. A. Mittermeier, and E. E. Louis. 2015. A New Species in the Genus Cheirogaleus (Cheirogaleidae). Primate Conserv. 29:43–54.

Li, H. 2009. SNPable Regions.

Li, H., and R. Durbin. 2009. Fast and accurate short read alignment with Burrows-Wheeler transform. Bioinformatics 25:1754–1760.

Li, H., and R. Durbin. 2011. Inference of human population history from individual whole-genome sequences. Nature 475:493–496. Nature Publishing Group.

Li, H., B. Handsaker, A. Wysoker, T. Fennell, J. Ruan, N. Homer, G. Marth, G. Abecasis, R. Durbin, and 1000 Genome Project Data Processing Subgroup. 2009. The Sequence Alignment/Map format and SAMtools. Bioinformatics 25:2078–2079.

Malaspinas, A., M. C. Westaway, C. Muller, V. C. Sousa, O. Lao, I. Alves, A. Bergström, G. Athanasiadis, J. Y. Cheng, J. E. Crawford, T. H. Heupink, E. Macholdt, S. Peischl, S. Rasmussen, S. Schiffels, S. Subramanian, J. L. Wright, A. Albrechtsen, C. Barbieri, I. Dupanloup, A. Eriksson, A. Margaryan, I. Moltke, I. Pugach, J. V. Moreno-mayar, S. Ni, F. Racimo, M. Sikora, T. S. Korneliussen, P. Ivan, S. Ellingvåg, G. Fourmile, P. Gerbault, D. Injie, G. Koki, M. Leavesley, B. Logan, A. Lynch, E. A. Matisoo-smith, P. J. Mcallister, A. J. Mentzer, M. Metspalu, A. B. Migliano, L. Murgha, M. E. Phipps, W. Pomat, D. Reynolds, F. Ricaut, P. Siba, M. G. Thomas, T. Wales, C. Ma, S. J. Oppenheimer, C. Tyler-smith, R. Durbin, J. Dortch, A. Manica, M. H. Schierup, R. A. Foley, M. M. Lahr, C. Bowern, J. D. Wall, T. Mailund, M. Stoneking, R. Nielsen, M. S. Sandhu, and L. Excoffier. 2016. A genomic history of Aboriginal Australia. Nat. Publ. Gr. 538:207–214. Nature Publishing Group.

Marçais, G., and C. Kingsford. 2011. A fast, lock-free approach for efficient parallel counting of occurrences of k-mers. Bioinformatics 27:764–770.

Martin, R.. 1972. Adaptive radiation and behaviour of the Malagasy lemurs. Proc. R. Soc. B Biol. Sci. 264.

Martin, S. H., J. W. Davey, and C. D. Jiggins. 2015. Evaluating the use of ABBA-BABA statistics to locate introgressed loci. Mol. Biol. Evol. 32:244–257.

Mckenna, A., M. Hanna, E. Banks, A. Sivachenko, K. Cibulskis, A. Kernytsky, K. Garimella, D. Altshuler, S. Gabriel, M. Daly, and M. A. Depristo. 2010. The Genome Analysis Toolkit: A MapReduce framework for analyzing next-generation DNA sequencing data. 1297–1303.

Mclain, A. T., R. Lei, C. L. Frasier, J. M. Taylor, C. A. Bailey, B. A. D. Robertson, S. D. Nash, J. C. Randriamanana, R. A. Mittermeier, and E. E. Louis. 2017. A New Cheirogaleus (Cheirogaleidae: Cheirogaleus crossleyi Group) Species from Southeastern Madagascar. Primate Conserv. 2017:27–36.

Meyer, W. K., A. Venkat, A. R. Kermany, B. Van De Geijn, S. Zhang, and M. Przeworski. 2015. Evolutionary history inferred from the de novo assembly of a nonmodel organism, the blue-eyed black lemur. Mol. Ecol. 24:4392–4405.

Morin, P., and K. B. Storey. 2009. Mammalian hibernation: Differential gene expression and novel application of epigenetic controls. Int. J. Dev. Biol. 53:433–442.

Nguyen, L. T., H. A. Schmidt, A. Von Haeseler, and B. Q. Minh. 2015. IQ-TREE: A fast and effective stochastic algorithm for estimating maximum-likelihood phylogenies. Mol. Biol. Evol. 32:268–274.

Nunziata, S. O., and D. W. Weisrock. 2018. Estimation of contemporary effective population size and population declines using RAD sequence data. Heredity (Edinb). 120:196–207.

Palkopoulou, E., S. Mallick, P. Skoglund, J. Enk, N. Rohland, H. Li, A. Omrak, S. Vartanyan, H. Poinar, A. Götherström, D. Reich, and L. Dalén. 2015. Complete genomes reveal signatures of demographic and genetic declines in the woolly mammoth. Curr. Biol. 25:1395–1400.

Pastorini, J., R. D. Martin, P. Ehresmann, E. Zimmermann, and M. R. Forstner. 2001. Molecular phylogeny of the lemur family cheirogaleidae (primates) based on mitochondrial DNA sequences. Mol. Phylogenet. Evol. 19:45–56.

Pauls, S. U., C. Nowak, M. Bálint, and M. Pfenninger. 2013. The impact of global climate change on genetic diversity within populations and species. Mol. Ecol. 22:925–946.

Prado-martinez, J., P. H. Sudmant, J. M. Kidd, H. Li, J. L. Kelley, B. Lorente-galdos, K. R. Veeramah, A. E. Woerner, T. D. O. Connor, G. Santpere, A. Cagan, C. Theunert, F. Casals, H. Laayouni, K. Munch, A. Hobolth, A. E. Halager, M. Malig, M. Pybus, L. Johnstone, M. Lachmann, J. Hernandez-rodriguez, I. Hernando-herraez, and K. Pru. 2013. Great ape genetic diversity and population history. Nature 499:471–475.

Putnam, N. H., B. L. O. Connell, J. C. Stites, B. J. Rice, M. Blanchette, R. Calef, C. J. Troll, A. Fields, P. D. Hartley, C. W. Sugnet, D. Haussler, D. S. Rokhsar, and R. E. Green. 2016. Chromosome-scale shotgun assembly using an in vitro method for long-range linkage. Genome Res. 26:342–350.

Quinlan, A. R., and I. M. Hall. 2010. BEDTools: A flexible suite of utilities for comparing genomic features. Bioinformatics 26:841–842.

Rakotoarinivo, M., A. Blach-overgaard, W. J. Baker, J. Dransfield, J. Moat, and J. Svenning. 2013. Palaeo-precipitation is a major determinant of palm species richness patterns across Madagascar: a tropical biodiversity hotspot Palaeo-precipitation is a major determinant of palm species richness patterns across Madagascar: a tropical biodiversity hot. Proc. Biol. Sci. 280.

Rogers, R. L., and M. Slatkin. 2017. Excess of genomic defects in a woolly mammoth on Wrangel island. PLoS Genet. 13:1–16.

Rumpler, Y., S. Crovella, and D. Montagnon. 1994. Systematic relationships among Cheirogaleidae (Primates, Strpsirhini) determined from analysis of highly repeated DNA. Folia Primatol 63:149–155.

Runemark, A., C. N. Trier, F. Eroukhmanoff, J. S. Hermansen, M. Matschiner, M. Ravinet, T. O. Elgvin, and G. P. Sætre. 2018. Variation and constraints in hybrid genome formation. Nat. Ecol. Evol. 2:549–556. Springer US.

Salmona, J., R. Heller, E. Quéméré, and L. Chikhi. 2017. Climate change and human colonization triggered habitat loss and fragmentation in Madagascar. Mol. Ecol. 26:5203–5222.

Sankararaman, S., S. Mallick, N. Patterson, and D. Reich. 2016. The Combined Landscape of Denisovan and Neanderthal Ancestry in Present-Day Humans. Curr. Biol. 26:1241–1247.

Schiffels, S., and R. Durbin. 2014. Inferring human population size and separation history from multiple genome sequences. Nat. Publ. Gr. 46:919–925. Nature Publishing Group.

Slon, V., F. Mafessoni, B. Vernot, C. de Filippo, S. Grote, B. Viola, M. Hajdinjak, S. Peyrégne, S. Nagel, S. Brown, K. Douka, T. Higham, M. B. Kozlikin, M. V. Shunkov, A. P. Derevianko, J. Kelso, M. Meyer, K. Prüfer, and S. Pääbo. 2018. The genome of the offspring of a Neanderthal mother and a Denisovan father. Nature 561:113–116. Springer US.

Song, S., E. Sliwerska, S. Emery, and J. M. Kidd. 2017. Modeling human population separation history using physically phased genomes. Genetics 205:385–395.

Soscia, S. J., and M. E. Harrington. 2005. Neuropeptide Y does not reset the circadian clock in NPY Y2−/− mice. Neurosci. Lett. 373:175–178.

Sun, H., J. Ding, M. Piednoe l, and K. Schneeberger. 2018. Genome analysis findGSE: estimating genome size variation within human and Arabidopsis using k-mer frequencies. Bioinformatics 34:550–557.

Team, R. C. 2015. R: A language and environment for statistical computing. R Found. Stat. Comput. Vienna, Austria.

Thiele, D., E. Razafimahatratra, and A. Hapke. 2013. Discrepant partitioning of genetic diversity in mouse lemurs and dwarf lemurs - Biological reality or taxonomic bias? Mol. Phylogenet. Evol. 69:593–609. Elsevier Inc.

Van der Auwera, G. A., M. O. Carneiro, C. Hartl, R. Poplin, G. Angel, A. Levy-moonshine, K. Shakir, D. Roazen, J. Thibault, E. Banks, K. V Garimella, D. Altshuler, S. Gabriel, and M. A. Depristo. 2013. From FastQ Data to High-Confidence Variant Calls: The Genome Analysis Toolkit Best Practices Pipeline. Curr. Protoc. Bioinforma. 43:1–33.

Vijay, N., C. Park, J. Oh, S. Jin, E. Kern, H. W. Kim, J. Zhang, and J. Park. 2018. Population Genomic Analysis Reveals Contrasting Demographic Changes of Two Closely Related Dolphin Species in the Last Glacial. 35:2026–2033.

Ward, B., and C. Van Oosterhout. 2016. HYBRIDCHECK: software for the rapid detection, visualization and dating of recombinant regions in genome sequence data. Mol. Ecol. Resour. 16:534–539.

Waterhouse, R. M., M. Seppey, F. A. Sim, and P. Ioannidis. 2017. BUSCO Applications from Quality Assessments to Gene Prediction and Phylogenomics Letter Fast Track. 35:543–548.

Wenzel, J. J., W. E. Kaminski, A. Piehler, S. Heimerl, T. Langmann, and G. Schmitz. 2003. ABCA10, a novel cholesterol-regulated ABCA6-like ABC transporter. Biochem. Biophys. Res. Commun. 306:1089–1098.

Williams, C. T., M. Radonich, B. M. Barnes, and C. L. Buck. 2017. Seasonal loss and resumption of circadian rhythms in hibernating arctic ground squirrels. J. Comp. Physiol. B Biochem. Syst. Environ. Physiol. 187:693–703. Springer Berlin Heidelberg.

Wilmé, L., S. Goodman, and J. Ganzhorn. 2006. Biogeographic Evolution of Madagascar’s Microendemic Biota. Science (80-.). 312:1063–1065.

Wilmé, L., C. Schwitzer, P. Devillers, and R. C. Beudels-jamar. 2014. Speciation in Malagasy lemurs: a review of the cryptic diversity in genus Lepilemur. Biotechnol. Agron. Soc. Environ. 18:577–588.

Yi, M., H. Li, Z. Wu, J. Yan, Q. Liu, C. Ou, and M. Chen. 2018. A Promising Therapeutic Target for Metabolic Diseases: Neuropeptide y Receptors in Humans. Cell. Physiol. Biochem. 45:88–107.

Yoder, A. D., C. R. Campbell, M. B. Blanco, M. dos Reis, J. U. Ganzhorn, S. M. Goodman, K. E. Hunnicutt, P. A. Larsen, P. M. Kappeler, R. M. Rasoloarison, J. M. Ralison, D. L. Swofford, and D. W. Weisrock. 2016. Geogenetic patterns in mouse lemurs (genus *Microcebus*) reveal the ghosts of Madagascar’s forests past. Proc. Natl. Acad. Sci. 113:8049–8056.

Yoder, A. D., J. Poelstra, G. P. Tiley, and R. Williams. 2018. Neutral Theory is the Foundation of Conservation Genetics. Mol. Biol. Evol. 35:1–5.

Young, M. D., M. J. Wakefield, G. K. Smyth, and A. Oshlack. 2010. Gene ontology analysis for RNA-seq: accounting for selection bias. Genome Biol. 11.

Yulyaningsih, E., L. Zhang, H. Herzog, and A. Sainsbury. 2011. NPY receptors as potential targets for anti-obesity drug development. Br. J. Pharmacol. 163:1170–1202.

Zamani, N., P. Russell, H. Lantz, M. P. Hoeppner, J. R. S. Meadows, N. Vijay, E. Mauceli, F. di Palma, K. Lindblad-Toh, P. Jern, and M. G. Grabherr. 2013. Unsupervised genome-wide recognition of local relationship patterns. BMC Genomics 14.

